# Cytoplasmic pool of spliceosome protein SNRNP70 regulates the axonal transcriptome and development of motor connectivity

**DOI:** 10.1101/2020.05.25.097444

**Authors:** Nikolas Nikolaou, Patricia M. Gordon, Fursham Hamid, Richard Taylor, Eugene V. Makeyev, Corinne Houart

## Abstract

Regulation of pre-mRNA splicing and polyadenylation plays a profound role in neurons by diversifying the proteome and modulating gene expression during development and in response to physiological cues. Although most pre-mRNA processing reactions are thought to occur in the nucleus, numerous splicing regulators are also found in neurites. Here, we show that U1-70K/SNRNP70, a major spliceosomal component, localizes in RNA-associated granules in axons. We identify the cytoplasmic pool of SNRNP70 as an important local regulator of motor axonal growth, nerve-dependent acetylcholine receptor (AChR) clustering and neuromuscular synaptogenesis. This cytoplasmic pool has a protective role for a limited number of axonal transcripts preventing them from degradation. Moreover, non-nuclear SNRNP70 is able to locally regulates splice variants of transcripts such as *agrin*, thereby locally controlling formation of synapses. Our results point to an unexpected, yet essential, function of local SNRNP70 in axonal development and indicate a role of splicing factors in local RNA metabolism during establishment and maintenance of neuronal connectivity.

## INTRODUCTION

RNA processing, transport and local translation play a central part in normal development and function of neural circuits. These processes are extensively regulated by numerous RNA-binding proteins (RBPs) interacting with RNAs inside and outside the nucleus (Darnell and Richter, 2012; Darnell, 2013). In recent years, much progress has been made in identifying the players and their roles in these processes, revealing a staggering complexity in the mechanisms of RNA metabolism and transport (Hocine, Singer and Grünwald, 2010; Turner-Bridger *et al*., 2018; Das, Singer and Yoon, 2019). Significantly, defects in RBP expression, localization and function have been linked with numerous neurodevelopmental and neurodegenerative diseases including autism, schizophrenia, bipolar disorders, spinal muscular atrophy, and amyotrophic lateral sclerosis (Bryant and Yazdani, 2016; Lee *et al*., 2016; Ito, Hatano and Suzuki, 2017; Khalil *et al*., 2018).

Many RBPs participate in the regulation of pre-mRNA splicing in the nucleus but some of these splicing regulators are also found in neurites of developing and matured neurons (Racca *et al*., 2010; Poulopoulos *et al*., 2019) For instance, the nucleus-rich RBP SFPQ additionally localizes to mouse dorsal root ganglion axons where it is thought to regulate RNA granule co-assembly and trafficking of mRNAs essential for axonal viability such as LaminB2 and Bclw (Cosker *et al*., 2016). SFPQ was also found to have an important function in axons of zebrafish motor neurons, contributing to normal development of this type of nerve cell (Thomas-Jinu *et al*., 2017). Pertinent to this study, we have previously detected the U1 small nuclear ribonucleoprotein 70K (SNRNP70), a core component of the spliceosome machinery, in zebrafish motor axons (Thomas-Jinu *et al*., 2017). These findings point to unexpected, and yet unknown, function for these proteins in axons.

SNRNP70 complexes with many other auxiliary proteins and U1 small nuclear RNA (U1 snRNA) to form the U1 snRNP component of the major spliceosome. SNRNP70 contains a tail domain mediating binding with the Sm proteins, an α-helix domain and an RNA recognition motif (RRM) forming contacts with the U1 snRNA and two low complexity arginine/serine domains involved in interactions with SR proteins (Pomeranz Krummel *et al*., 2009; Kondo *et al*., 2015). SNRNP70 is required for both constitutive and alternative pre-mRNA splicing (Wahl, Will and Lührmann, 2009). Its function outside the nucleus is currently unknown.

Here, we show that SNRNP70 localizes to cytoplasmic puncta closely associated with RNA granules within axonal compartments. We demonstrate that the cytoplasmic pool of SNRNP70 is an important regulator of motor axonal growth, nerve-dependent acetylcholine receptor (AChR) clustering and neuromuscular synaptogenesis. We present evidence supporting a SNRNP70-mediated regulation of local mRNA processing. Together, our study identifies a novel function of a major spliceosomal protein in neurites, modulating the local transcriptome during circuit formation.

## RESULTS

### SNRNP70 co-localizes with RNAs in developing axons

Using an antibody against a highly conserved region of the human SNRNP70 protein, we examined SNRNP70 protein distribution in zebrafish. In line with the key role of SNRNP70 in pre-mRNA splicing, we found robust nuclear staining across embryonic and larval tissues including GFP-labeled motor neurons in *Tg(hb9:GFP)* animals (Figure S1A and data not shown). As previously observed (Thomas-Jinu *et al*., 2017), SNRNP70 was also detected inside motor axons innervating the muscles at all stages examined (Figure S1A). In addition, SNRNP70 immunoreactivity could be seen within motor axon terminals adjacent to AChRs clusters (Figure S1B), indicating that SNRNP70 also localizes to sites proximal to neuromuscular junctions (NMJs), synapses formed between motor neurons and muscle fibers. To investigate the SNRNP70 protein distribution at a subcellular resolution, we dissociated and cultured neurons derived from *Tg(hb9:GFP)* zebrafish embryos at 28 hours post fertilization (hpf). At one day *in vitro* GFP^+^ neurons had visible growth cones extending from the soma and in some cases formed simple axonal arbours (Figure 1A). Immunostaining showed that SNRNP70 protein localized to axons of these GFP^+^ neurons in the form of small puncta (Figure 1A).

**Figure 1:**
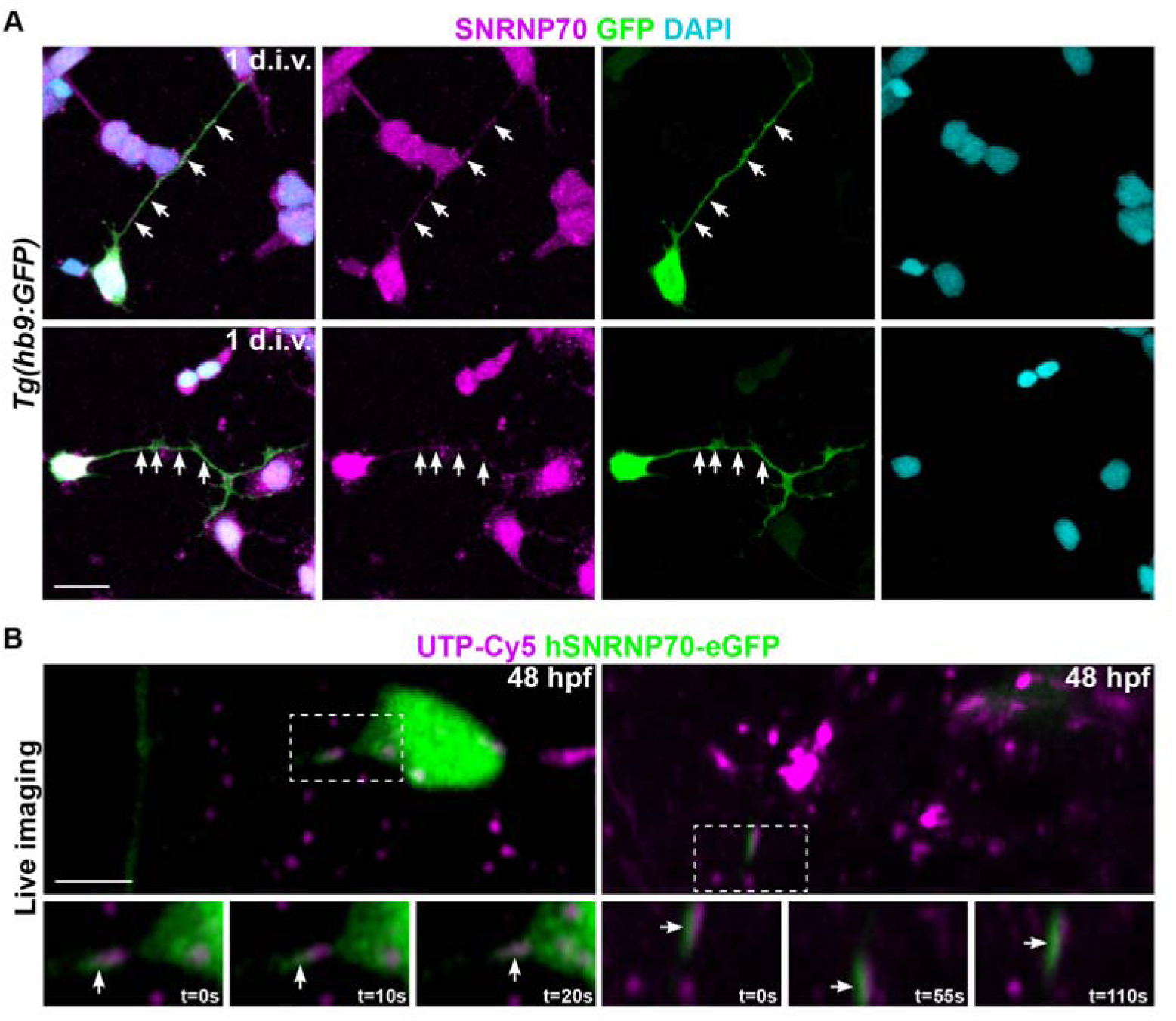
SNRNP70 localizes in axons. **(A)** Single plane confocal images of dissociated motor neurons at one day in vitro derived from *Tg(hb9:GFP)* animals at 28 hpf. Neurons were stained with anti-SNRNP70 to reveal the localization of the protein in neurites and DAPI to mark nuclei. Arrows indicate single SNRNP70-positive puncta within GFP^+^ axons. Representative images from two independent experiments. Scale bar, 10 μm. **(B)** Time-lapse of larval neurons at 48 hpf sparsely labeled with hSNRNP70-eGFP. Top: images show GFP labeled interneurons co-expressing UTP-Cy5. Bottom: insets depict axon segments showing association between Cy5-RNA granules (magenta) and hSNRNP70-eGFP (green) signals (arrows indicate the same punctum during the course of the imaging session with time shown in the bottom right corner of individual time points). Representative examples from three independent experiments. Scale bar, 5 μm. See also Figure S1 and Movies S1 and S2.

Several RNA binding proteins are known to associate with ribonucleoprotein particles (RNPs) transporting RNA molecules along axons and dendrites (Ortiz *et al*., 2017; Chu *et al*., 2019). To investigate this possibility for the cytoplasmic fraction of SNRNP70 we imaged a functionally active eGFP-tagged version of human SNRNP70 (hSNRNP70-eGFP) (Huranová *et al*., 2010). We also labeled newly synthesized RNAs with a fluorescent derivative of uridine-5’-triphosphate (Cy5-UTP) to allow subsequent visualization of fluorescent Cy5-RNA^+^ RNP granules in axons (Wong *et al*., 2017). Confirming our immunofluorescence data above, mosaic expression of the recombinant hSNRNP70-eGFP protein showed robust nuclear localization in hindbrain and spinal cord interneurons (Figure 1B). Importantly, we also observed multiple eGFP puncta co-localizing with RNAs in neuronal processes. Some of these puncta appeared to be oscillatory whereas others were motile (Figure 1B, Videos S1 and S2). These results suggest that SNRNP70 is present in axons where it associates and moves with RNAs.

### SNRNP70 is cell-autonomously required in motor neurons for neuromuscular assembly

As small deletions preventing the cytosolic localisation would also affect the protein function, we assessed the cytoplasmic function of SNRNP70 by generating a complete loss of function allele in which we integrated an inducible, cytoplasmic-localized SNRNP70 transgenic line to drive expression in the null zebrafish animal. The null allele was generated by targeting exon2, the first coding exon of the zebrafish *snrnp70* gene, using appropriate CRISPR/Cas9 reagents. DNA sequencing of target-specific PCR products confirmed that the *snrnp70*-targeted allele (*kg163* allele) carried a deletion of 13 bases (Figure S2A). The deletion resulted in a frameshift immediately after amino-acid 16 and a truncation 66 amino-acids after the mutation (Figures S2B and S2C). The mutation was predicted to disrupt all known functional domains of the SNRNP70 protein, and indeed the antibody did not detect any protein in the null (Figure S2D). In addition to truncating the open reading frame, our data show that the mutant mRNA transcript was likely targeted by nonsense-mediated decay (NMD; data not shown) which is expected for RNA with a premature translation termination codon. Homozygous mutants for *snrnp70* (*snrnp70*^*−/−*^ thereafter referred as null) were obtained from heterozygotes crosses.

Due to maternal RNA contribution, early development of *snrnp70*-null embryos was not affected until mid-somite stage (data not shown). From 18-somite stage onwards, we observed morphological defects including abnormal tail extension, heart edema and cell death. Brain regions were particularly sensitive to the loss of *snrnp70* indicating a critical role of nuclear SNRNP70 in neural development (Figure S2E). Despite the dramatic defects seen in the brain, the spinal cord and somites were much less affected between 18-28 hpf allowing us to examine the role of cytoplasmic SNRNP70 in the formation of the NMJ, a synapse established early in neural circuit development. In zebrafish there are four subtypes of primary MNs and several subtypes of secondary MNs. Primary MN growth cones exit the spinal cord around 18 hpf and reach the most ventral part of the myotome by 28 hpf (Myers, Eisen and Westerfield, 1986). Using the *Tg(hb9:GFP)* line we examined how the absence of the SNRNP70 protein affected normal MN development. The gross morphology of null MNs was indistinguishable from siblings at 28 hpf, apart from a reduction in the total length and thickness of the motor nerves (Figures 2A-2C). At 48 hpf, motor axonal development was severely disrupted; several aberrant branches, abnormal branching within the myotome and reduced innervation of the lateral myosepta were observed (Figures S3A-S3C).

**Figure 2:**
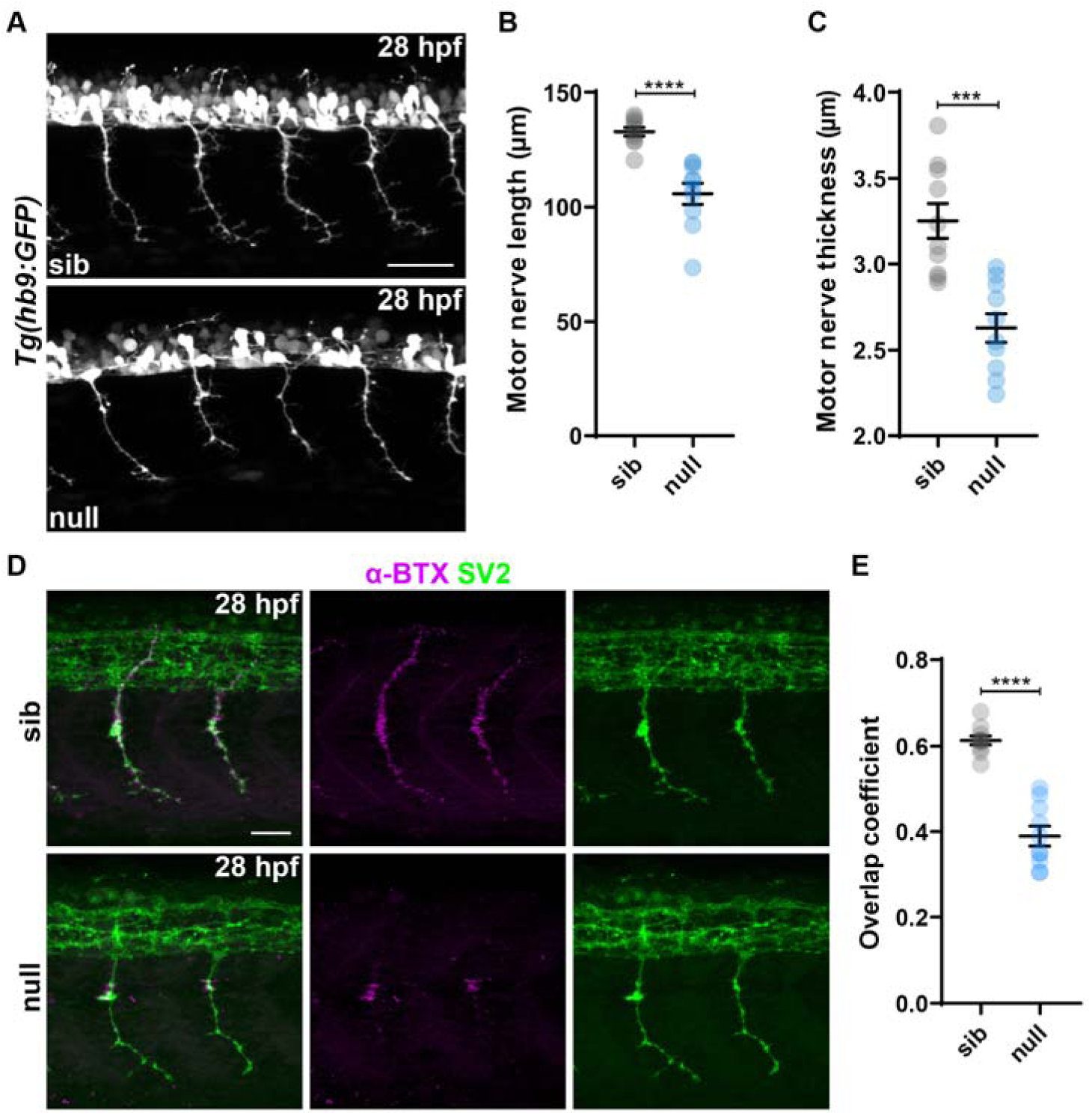
Loss of SNRNP70 affects motor neuron growth and neuromuscular connectivity. **(A)** Representative confocal images of *Tg(hb9:GFP)* sibling and null embryos at 28 hpf showing motor nerves innervating the myotome. Scale bar, 50 μm. **(B-C)** Quantifications showing the length and thickness of motor nerves in the two groups. All graphs show mean values ± SEM. **** *P* < 0.0001; *** *P* < 0.001, two-tailed unpaired t-test, *n* = 10 animals per group in two independent experiments. **(D)** Confocal images of sibling and null embryos at 28 hpf stained with anti-SV2 antibody to mark the pre-synaptic locations and a fluorescently tagged α-BTX to label post-synaptic structures. Scale bar, 25 **(E)** Quantification of the degree of overlap between SV2 and α-BTX. The graph shows mean values ± SEM. **** *P* < 0.0001, two-tailed unpaired t-test, *n* = 10 animals per group in two independent experiments. See also Figures S2 and S3.

One of the earliest hallmarks of the NMJ assembly is the clustering of AChRs on the surface of muscle cells adjacent to the muscle-innervating axons, replacing the initial aneural AChR clustering (Flanagan-Steet *et al*., 2005; Panzer, 2006; Wu, Xiong and Mei, 2010). To investigate how neuromuscular assembly may be affected in *snrnp70*-null embryos, we stained pre-synaptic MN terminals with an SV2-specific antibody and AChR-containing post-synaptic structures on muscle fibers with a fluorescently tagged α-bungarotoxin (α-BTX). At 28 hpf there were visible clusters of AChRs apposed to the SV2^+^ pre-synaptic sites in siblings. However, in null animals the AChR clusters were virtually absent while the SV2^+^ structures appeared normal (Figure 2D). Consistent with this phenotype, there was a significant decrease in the degree of co-localization between SV2 and α-BTX within the myotome (Figure 2E). Reduced AChR clustering was also evident at 48 hpf, a stage of active neuromuscular assembly (Figures S3D and S3E). The lack of post-synaptic AChRs was unlikely due to abnormal development of the myotome. Indeed, somitic muscle differentiation occurs normally in the null embryos, with typical V-shaped somitic boundaries, proper differentiation of both slow and fast muscle fibres, and formation of sarcomeres (Figures S3F and S3G). The absence of AChR clusters on muscle fibres, without initial defects in muscle differentiation, indicates that the primary defect is in the MNs.

To assess whether SNRNP70 is required cell-autonomously in MNs for the induction of AChRs at NMJs, we performed cell mosaic experiments. Cell transplantations from *Tg(hb9:GFP)* donors into the spinal neural plate territory of wild-type hosts at 70% epiboly resulted in clones of 1-4 fluorescently marked MNs. Sibling MNs transplanted into WT embryos produced axonal projections fully integrated into the host environment. Quantification of the appositions formed by transplanted neurons with host muscle fibers showed that NMJs formed between the sibling MNs and the WT host environment are not only evident within the axonal arbour, but also along the main axonal shaft of the transplanted MNs (Figure 3A). Transplantations of null MNs into WT hosts resulted in normal ventral projections into the myotome at 48 hpf, but absent or much reduced clustering of AChRs within the WT host environment, at the level of both axonal arbours and the axonal shaft (Figure 3B). These observations show that SNRNP70 is required cell-autonomously in MNs for the clustering of AChR at the NMJ.

**Figure 3:**
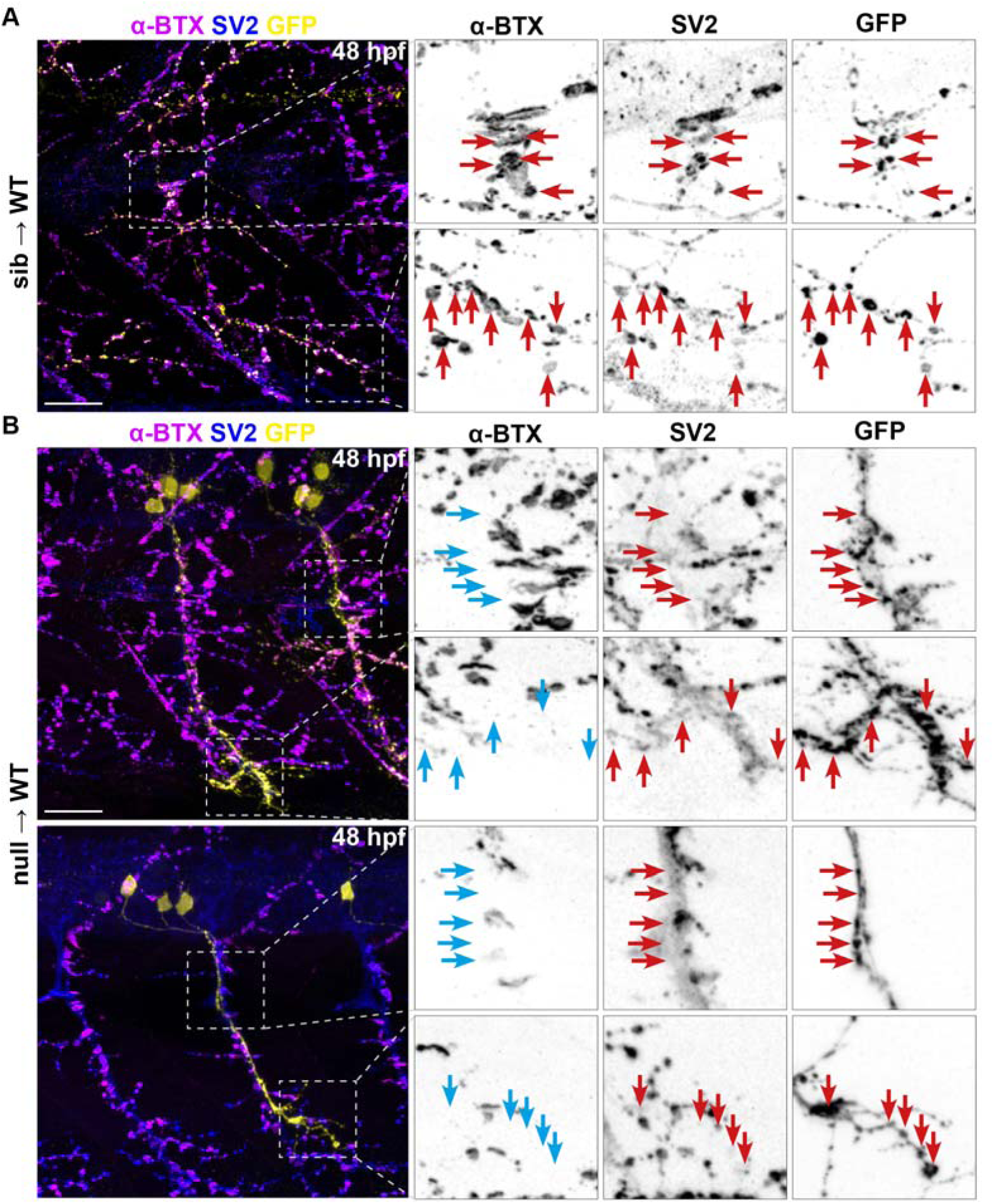
SNRNP70 is cell-autonomously required in motor neurons for neuromuscular assembly. **(A)** Confocal image showing transplanted sibling motor neurons derived from a *Tg(hb9:GFP)* line within a WT host animal that has been stained for anti-SV2 and α-BTX to mark pre- and post-synaptic sites, respectively. Right: insets from two regions in the image on the left depicting the apposition of pre- and post-synaptic sites (red arrows) within the GFP^+^ motor neurons. Representative images from five independent experiments. Scale bar, 25 μm. **(B)** Examples of GFP^+^ transplanted null motor neurons within WT host animals that have been stained with anti-SV2 and α-BTX. Right: insets from regions in the images on the left showing absence of α-BTX^+^ puncta (cyan arrows) juxtaposed to SV2 labelling within the GFP^+^ null motor neurons (red arrows). Representative images from three independent experiments. Scale bar, 25 μm.

### The cytoplasmic pool of SNRNP70 is sufficient for neuromuscular connectivity

We wondered if the newly identified role of SNRNP70 in NMJ development depended on the cytoplasmic fraction of this RBP. As stated above, to answer this question, we generated a transgenic line of zebrafish expressing hSNRNP70-eGFP from a UAS promoter – *Tg(UAS:hSNRNP70-eGFP)*. Crossing these fish to the inducible *Tg(ubi:ERT2-Gal4)* line to obtain double transgenic embryos allows ubiquitous expression of hSNRNP70-eGFP upon the addition of 4-OHT (Figure S4A). We analysed four groups of embryos obtained from heterozygote *snrnp70*^*kg163*^ transgenic crosses: (1) siblings that are GFP^−^ (sib/GFP^−^); (2) siblings that are GFP^+^ (sib/GFP^+^); (3) nulls that are GFP^−^ (null/GFP^−^); and (4) nulls that are GFP^+^ (null/GFP^+^). Transgenic expression of full-length hSNRNP70-eGFP in siblings did not interfere with the normal development of these animals (Figure S4C), indicating that the additional SNRNP70 protein is not toxic to the cells. Moreover, expression of hSNRNP70-eGFP in null embryos rescued the morphological abnormalities and restored motor function and neuromuscular assembly (Figures S4C-S4F).

To generate a cytoplasmic form of hSNRNP70, we deleted the two nuclear localization signals (NLS) using the mutagenesis strategy previously shown to increase cytoplasmic localization of the hSNRNP70 without affecting its structure and/or functional domains (Romac, Graff and Keene, 1994). In addition, we inserted nuclear export signals (NES) at the C-terminal end of the NLS-deficient protein to ensure that any nuclear hSNRNP70 is rapidly exported to the cytoplasm. We found that three copies of NES were sufficient to completely restrict hSNRNP70-eGFP localization to the cytoplasm. Using this construct we generated a second UAS-transgenic line – *Tg(UAS:hSNRNP70ΔNLS3xNES-eGFP)* – which when crossed with the *Tg(ubi:ERT2-Gal4)* line allows ubiquitous expression of cytosolic hSNRNP70 (Figure S4B). We found that transgenic expression of cytosolic hSNRNP70-eGFP in sibling animals did not substantially affect their normal development (Figure 4A), indicating that this cytoplasmic form of SNRNP70 does not interfere with the normal function of SNRNP70 in the nucleus. Expression of cytosolic hSNRNP70-eGFP in null mutants was unable to rescue their morphological defects (Figure 4A), however, there was a restoration in touch-evoked startle response (Figure 4B), an increase in AChR clustering (Figures 4C and 4D) and overall growth of motor nerves (Figure 4E). These experiments demonstrate that the cytoplasmic pool is sufficient to drive SNRNP70 function required for motor axonal growth and synaptogenesis.

**Figure 4:**
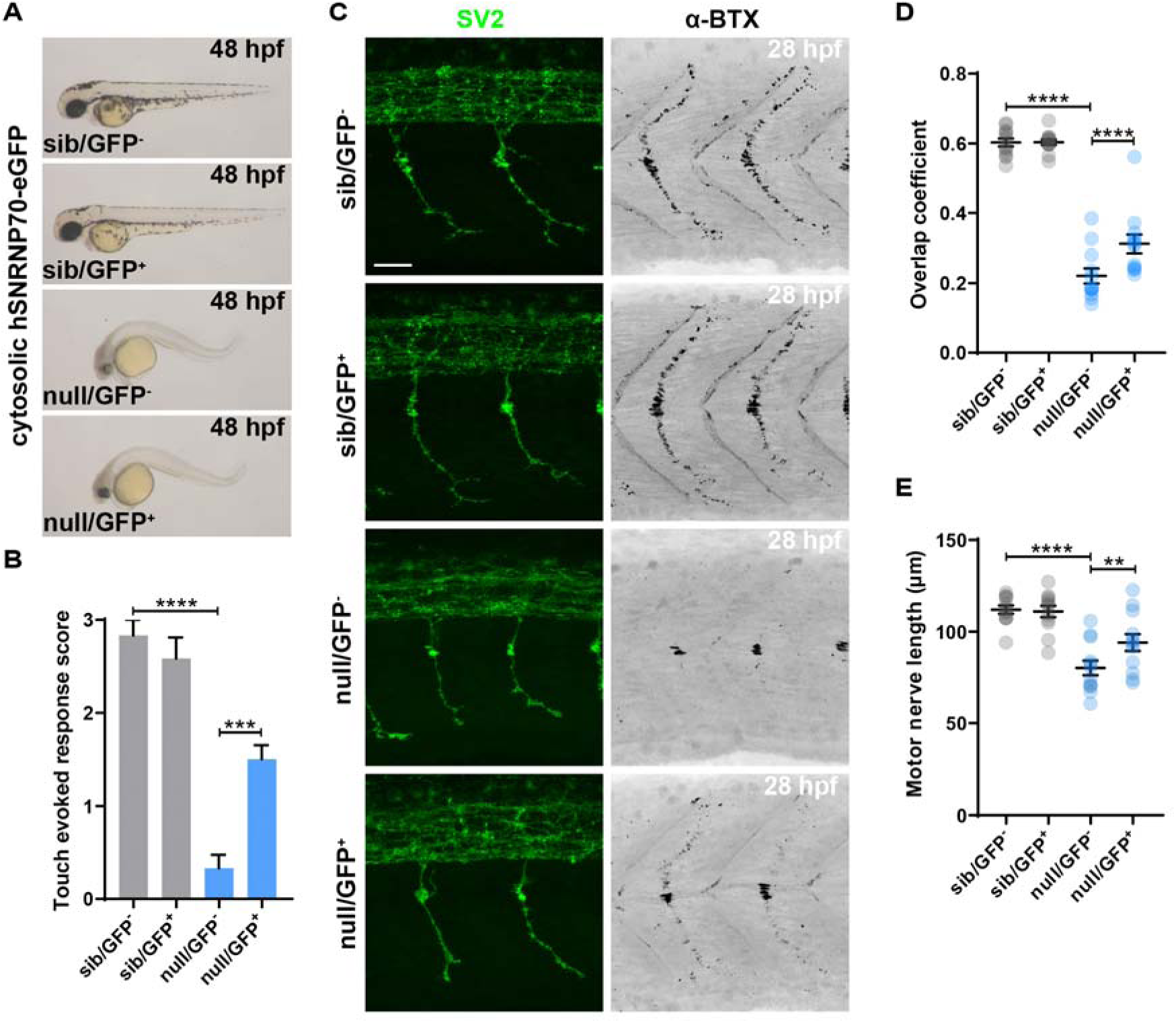
Overexpression of cytosolic hSNRNP70-eGFP rescues motor neuron growth and neuromuscular assembly. Overexpression of cytosolic hSNRNP70-eGFP in embryos was achieved using the *Tg(ubi:ERT2-Gal4;UAS:hSNRNP70ΔNLS3xNES-eGFP)* line. Embryos derived from the same clutch were sorted into four groups according to genotype: (1) sib/GFP^−^, (2) sib/GFP^+^, (3) null/GFP^−^ and (4) null/GFP^+^. **(A)** Representative images of embryos at 48 hpf from three independent experiments. **(B)** Quantification of touch-evoked startle response at 48 hpf. The graph shows mean values ± SEM. **** *P* < 0.0001; *** *P* < 0.001, One-way ANOVA, *n* = 12 animals per group in three independent experiments. **(C)** Confocal images of embryos at 28 hpf stained with anti-SV2 antibody to mark the pre-synaptic locations and a fluorescently tagged α-BTX to label post-synaptic structures. Representative images from three independent experiments. Scale bar, 25 μm. **(D)** Quantification of the degree of overlap between SV2 and α-BTX at 28 hpf in the four groups. The graph shows mean values ± SEM. **** *P* < 0.0001, One-way ANOVA, *n* = 12 animals per group in three independent experiments. **(E)** Quantification of the length of motor nerves at 28 hpf. The graph shows mean values ± SEM. **** *P* < 0.0001; ** *P* < 0.01, One-way ANOVA, *n* = 12 animals per group in three independent experiments. See also Figure S4.

### Cytoplasmic SNRNP70 modulates gene expression and alternative splicing

To assess whether cytoplasmic SNRNP70 has an effect upon the local transcriptome, we analysed 28 hpf sibling and null embryos either positive or negative for cytosolic hSNRNP70-eGFP by RNA sequencing (RNA-seq). A principal component analysis of the gene expression data showed a clear segregation between sibling and null animals regardless of the expression of cytosolic hSNRNP70-eGFP (Figure S5A). We first evaluated gene expression changes across all annotated genes in zebrafish and discovered 2015 genes upregulated and 2498 genes downregulated following the loss of SNRNP70 (Figure S5B and Table S1).

To assess whether cytoplasmic SNRNP70 was sufficient to restore the expression of some transcripts to wild-type-like levels in null embryos we compared the null/GFP^+^ to the null/GFP^−^ group. We found 347 genes (from a total of 1009 genes whose expression changed significantly between these two groups) corresponding to significant transcript rescue events (Figures S5C and S5D and Table S2). Gene Ontology (GO) enrichment analysis of the 347 restored transcript levels showed association, among others, to structural constituents of the ribosome as well as protein targeting to ER and plasma membrane affecting nervous system development in particular (Figure S5E). Taken together, these data suggest that the cytoplasmic fraction of SNRNP70 may regulate local transcripts abundance independently of its nuclear roles.

Given that SNRNP70 is an essential splicing factor, we also examined changes in pre-mRNA splicing patterns (Figure 5A). Loss of SNRNP70 (comparing null/GFP^−^ to sib/GFP^−^ group) resulted in significant alternative splicing changes with most altered events to be either cassette exon (CE) splicing or intron retention (IR) in null/GFP^−^ animals. Changes in alternative 3’ (Alt3) and 5’ (Alt5) usage were also observed (Figure 5B and Table S3). In the case of CE, the majority of altered events resulted in shorter alternatively spliced isoforms (i.e. increased exon skipping) in null/GFP^−^ animals, whereas in the case of IR there was an equal number of increases and decreases in IR resulting in longer and shorter alternative splicing isoforms, respectively, in null/GFP^−^ compared to control sib/GFP^−^ animals (Figure 5B).

**Figure 5:**
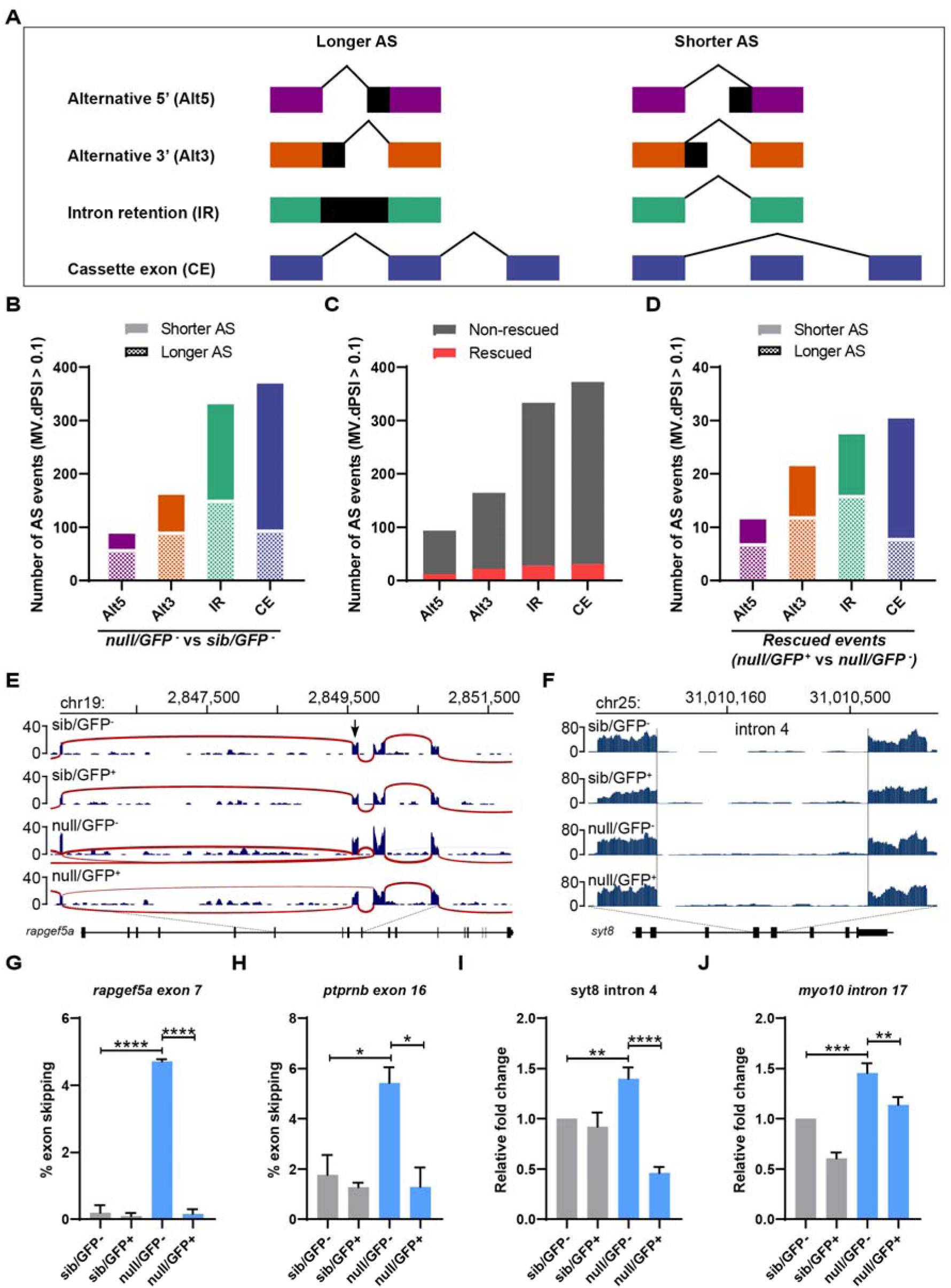
Loss of SNRNP70 leads to alternative splicing errors. Data are based on RNA-seq in 28 hpf embryos derived from overexpression of cytosolic hSNRNP70-eGFP in sibling and null animals using the *Tg(ubi:ERT2-Gal4;UAS:hSNRNP70ΔNLS3xNES-eGFP)* line. Embryos were sorted into four groups according to genotype: (1) sib/GFP^−^, (2) sib/GFP^+^, (3) null/GFP^−^ and (4) null/GFP^+^. **(A)** Key for alternative splicing changes shown in **(B-D)**. **(B)** Graph showing the number of alternative splicing events that appear significantly different (MV.dPSI > 0.1) comparing null/GFP^−^ to sib/GFP^−^ embryos. *n* = 3 biological samples per genotype. **(C)** Graph showing the number of alternative splicing events that are significantly (MV.dPSI > 0.1) restored (shown in red) or not (gray) following overexpression of cytosolic hSNRNP70-eGFP in null embryos. These data are based on comparison between sib/GFP^−^ vs null/GFP^−^ and then between null/GFP^−^ vs null/GFP^+^. **(D)** Graph of the 93 rescued alternative splicing events from **(C)** depicting the effect SNRNP70 loss had on these events prior to the rescue. These data are based on comparison between sib/GFP^−^ vs null/GFP^−^ and then between null/GFP^−^ vs null/GFP^+^. **(E-F)** Schematic plots for a representative example of CE (*rapgef5a exon7*) and IR (*syt8 intron 4*), respectively, in the four experimental groups. **(G-H)** Semiquantitative RT-PCR validation of exon skipping events (*rapgef5a exon7* and *ptprnb exon16*). The graphs show mean values ± SEM of the percentage of exon skipping over the total expression of that transcript in four experimental groups. **** *P* < 0.0001; ** *P* < 0.01; * *P* < 0.05. One-way ANOVA, two technical replicates in two independent biological samples. **(I-J)** qPCR validation of intron retention events (*syt8 intron 4* and *myo10 intron 17*). The graphs show mean values ± SEM of the relative expression of the intron retained transcript normalized to the total expression of that transcript in four experimental groups. **** *P* < 0.0001; *** *P* < 0.001; ** *P* < 0.01. One-way ANOVA, two technical replicates in two independent biological samples. See also Figures S5, S6 and S7.

Surprisingly, transgenic expression of cytosolic hSNRNP70-eGFP in nulls (comparing null/GFP^+^ to null/GFP^−^ group) recovered a small fraction of affected alternatively spliced events. These 93 rescued events were from all types of alternative splicing, with CE and IR being the most abundant categories (Figure 5C and Table S4). The majority of CE restored events were defective exon skipping events in null/GFP^−^ animals, whereas the majority of IR rescued events were increase of abnormal IR in null/GFP^−^ animals (Figure 5D). Schematic plots for representative examples of CE and IR events are highlighted in Figures 5E and 5F, respectively.

Interestingly, GO enrichment analysis of the rescued alternatively spliced events showed a significant enrichment for synaptic vesicle recycling proteins, protein tyrosine phosphatase signalling molecules and vesicle tethering complexes (Figure S6A), i.e. essential components of axonal growth and synapse function. Several cytoplasmic SNRNP70-dependent CE events were tested by semi-quantitative RT-PCR. We validated two CE splicing events (*rapgef5a* exon 7 and *ptprnb* exon 16). Supporting the RNA-seq results, we found exon skipping to be significantly increased in null/GFP^−^, and their splicing was significantly recovered in null/GFP^+^ (Figures 5G and 5H). We also tested several IR events by RT-qPCR and were able to validate two IR events (*syt8* intron 4 and *myo10* intron 17), showing significant increase in null/GFP^−^, and a significantly restored normal splicing in null/GFP^+^ (Figures 5I and 5J). These rescues are unlikely to be carried out in the nucleus by rescued splicing factors as the expression of all spliceosomal proteins (with the exception of SNRNP70) either remained unchanged or increased in null/GFP^−^ animals, which is then unaffected following expression of cytoplasmic SNRNP70 (Table S5). Altogether, these data suggest that the cytoplasmic fraction of SNRNP70 modulates the alternative splicing of transcripts associated to neuronal development and synaptic function.

### Cytoplasmic SNRNP70 protects transcripts from NMD through alternative splicing

Expression of cytoplasmic SNRNP70 was sufficient to restore the expression of a subset of affected genes towards control levels. We detected a somewhat stronger enrichment for genes downregulated in null animals and upregulated by cytoplasmic SNRNP70 (Figures S7A and S7B). This observation suggests that cytoplasmic SNRNP70 may protect some of these transcripts from degradation. Alternative splicing can alter gene expression levels by producing RNA isoforms containing premature termination codons and thus subjected to NMD. To test whether the gene downregulation effect caused by the loss of SNRNP70 can be at least in part due to this mechanism, we quantified the number of alternative splicing events within each group of genes whose expression is regulated by SNRNP70. We found that the incidence of splicing changes was significantly higher for the group of transcripts downregulated by SNRNP70 knockout compared to non-regulated genes. Downregulated transcripts rescued by cytoplasmic SNRNP70 showed an even stronger enrichment for alternative splicing (Figure S7C). Intriguingly, SNRNP70-rescued downregulated transcripts also had the highest percentage of significant alternative splicing events leading to NMD (Figure S7D). Moreover, the percentage of regulated splicing events predicting NMD is significantly higher among the SNRNP70-rescued alternative splicing events (Figure S7E and Table S4). Thus, the production of NMD-sensitive isoforms in the absence of SNRNP70 may explain a subset of regulation effects in the knockout and suggests a possible protection from NMD in presence of cytoplasmic SNRNP70.

### The cytoplasmic fraction of SNRNP70 controls the production of Z+AGRN isoforms

Given the ability of cytoplasmic SNRNP70 to modulate alternative splicing events, we wondered whether this modulation could explain the rescue of neuromuscular connectivity in null animals. Agrin (AGRN) is a multidomain proteoglycan secreted by the MN growth cone which binds to LRP4/Musk receptor complex on the muscle fibre plasma membrane. The activation of this receptor complex leads to a signal transduction pathway that induces the neural clustering of AChRs (Glass *et al*., 1996; Herbst, 2000; Kim *et al*., 2008; Zhang *et al*., 2008). It has been previously shown that the ability of AGRN to induce the clustering of AChR depends on alternative splicing to produce Z+AGRN isoforms that have an insertion of either or both of two micro-exons encoding eight (z8) or eleven (z11) amino acids, respectively (Ferns *et al*., 1993; Hoch *et al*., 1994; Gesemann, Denzer and Ruegg, 1995; Burgess *et al*., 1999).

Therefore, we set out to examine whether the alternative splicing of Z+AGRN isoforms is impaired in null embryos and if so, whether this was rescued by expression of cytosolic hSNRNP70-eGFP. Since the annotation of zebrafish genome is currently incomplete, these exons are absent from the database of alternative events that we used in our bioinformatics analysis. To address this problem, we took an RT-PCR-based approach. Through sequence alignment with mouse and human we mapped the two putative micro-exons (named z8 and z11) between exons 33 and 34 of the zebrafish AGRN locus. Three alternatively spliced *z+agrn* variants were identified, containing insertions of either eight (33-z8-34), eleven (33-z11-34) or nineteen amino acids (33-z8-z11-34), respectively. Moreover, comparison of the amino-acid sequence of Z+AGRN domain encoded by the two micro-exons showed sequence conservation between human and zebrafish (Figure 6A). Following RT-qPCR analysis, we found that the while the total transcript expression of non-*z+agrn* (*agrin 33-34*) isoforms increases (Figure 6B), the expression of the three *z+agrn* transcripts is reduced following the loss of SNRNP70 (Figures 6C-6E). Intriguingly, overexpression of cytosolic hSNRNP70-eGFP brought expression levels of non-*z+agrn* isoforms back to normal levels and rescued the expression of two out of three *z+agrn* isoforms (33-z8-34 and 33-z11-34) (Figures 6B-6E). These results indicate that (1) the cytoplasmic pool of SNRNP70 regulates the normal production of Z+AGRN isoforms, (2) this regulation involves reciprocal changes in fractional abundance of *z+* and non-*z+agrn* splice variants and (3) changes in the expression of *z+agrn* isoforms may partly underly the ability of cytoplasmic SNRNP70 to control neuromuscular connectivity.

**Figure 6:**
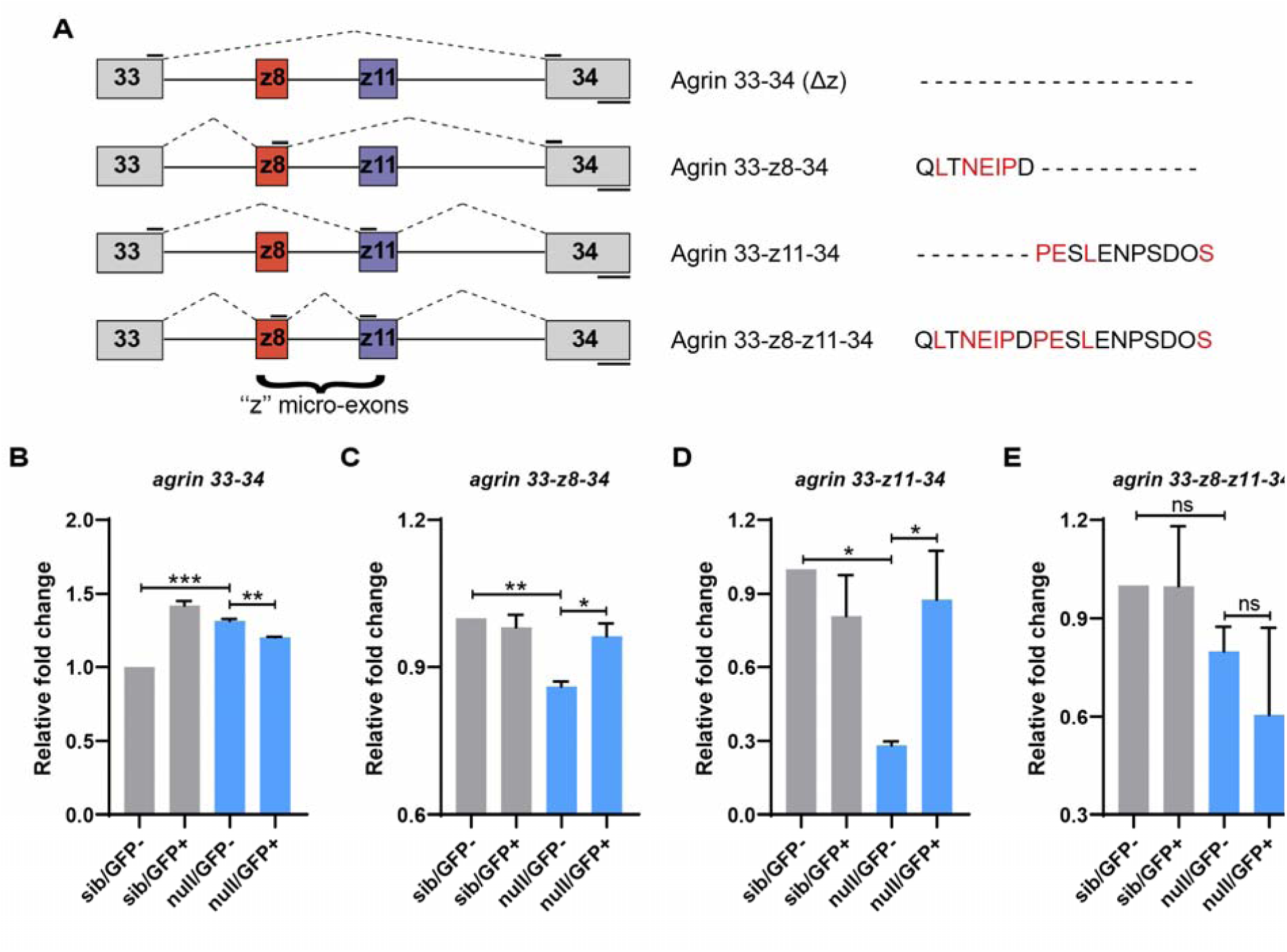
Cytoplasmic SNRNP70 regulates the alternative splicing of Z+AGRN isoforms. **(A)** Agrin Z+ isoforms identified by RT-PCR are shown schematically, with conserved residues between human and zebrafish shown in red. Lines depict the position of primers used for the qPCR analysis in **(B-E)**. **(B-E)** qPCR analysis in sibling and null animals as well as following overexpression of cytosolic hSNRNP70-eGFP using *Tg(ubi:ERT2-Gal4; UAS:hSNRNP70ΔNLS3xNES-eGFP)* embryos at 28 hpf. Primers were designed to specifically amplify the total expression of *agrin* (*agrin 33-34*), as well as the three *z+agrin* isoforms (*33-z8-34, 33-z11-34* and *33-z8-z11-34*). The graphs show mean values ± SEM for fold change in expression in relation to control animals (sib/GFP^−^). *** *P* < 0.001; ** *P* < 0.01; * *P* < 0.05; ns, not significant, One-way ANOVA, two technical replicates in two independent biological samples.

## DISCUSSION

The wiring of the nervous system is a complex process that depends on the availability of specific mRNAs and proteins at the right time (e.g. during axon guidance or synapse formation) and in the right place (e.g. growth cone, synaptic sites). Our study shows that the major spliceosomal protein SNRNP70 is closely associated with mRNA molecules within neurites and carries essential non-nuclear functions in neurons. Moreover, we identify cytoplasmic SNRNP70 as an important regulator of neuromuscular connectivity and show that this is achieved through an SNRNP70-mediated modulation of local pre-mRNA processing. We show that cytoplasmic SNRNP70 is functionally sufficient for the processing of a restricted set of neuronal transcripts during development, indicating a likely role for axonal SNRNP70 in establishment and maintenance of neuronal connections.

It is widely understood that U1 snRNP assembly initiates within the cytoplasm before SNRNP70 (independently of U1 snRNA and the Sm core of the particle) is transported to the nucleus (Fischer *et al*., 1993; Romac, Graff and Keene, 1994; Fischer, Liu and Dreyfuss, 1997; Liu *et al*., 1997). Our study provides the first evidence that the cytoplasmic pool of SNRNP70 is functionally active prior to its nuclear translocation. One possibility is that SNRNP70 may be involved in a form of extra-nuclear pre-mRNA processing. Tissue specific alternative splicing is a powerful source of protein diversity. This mechanism is particularly important in the nervous system where isoform variation is involved in cell recognition, synapse formation, synaptic plasticity, ion channel function and neurotransmission (Grabowski and Black, 2001; Norris and Calarco, 2012). While post-transcriptional regulation commonly occurs in the nucleus, recent studies indicate that spliceosomal components and splicing factors are found in axons and dendrites where they retain their potential to promote pre-mRNA splicing (Glanzer *et al*., 2005; König *et al*., 2007; Poulopoulos *et al*., 2019). This suggests that SNRNP70 could be interacting with other splicing factors to promote the complete splicing of partially spliced pre-mRNAs found in neurites. For instance, Nova1 and Nova2, two well conserved neuron-specific splicing factors, regulate the alternative splicing of *z+agrn* isoforms and are essential for Z+AGRN-mediated AChR cluster formation at the NMJ (Ruggiu *et al*., 2009). Nova proteins were also found to regulate brain- and subtype-specific alternative splicing programs required for axon guidance and neural circuit formation (Leggere *et al*., 2016; Saito *et al*., 2016). Like SNRNP70, Nova proteins were also found to localize in neurites (Racca *et al*., 2010). Therefore, it will be interesting to see whether extra-nuclear SNRNP70 and Nova proteins interact in the context of axon guidance and motor connectivity. The possibility for local alternative splicing is an attractive idea in the context of neurons. Many of the neuronal populations controlling our behaviours have the nucleus tens of centimeters away from the site of active synapses. The option of local splicing would be an advantageous mechanism to generate alternative transcripts known to be essential for the rapid modulation of synaptic connectivity and other neuronal functions.

Our studies also point to a role of NMD in SNRNP70-controlled gene regulation events. Alternative splicing can act not only to generate protein isoform diversity but also to quantitatively control gene expression levels within cells. It is known that several regulated alternative splicing events do not lead to generation of protein products, but instead they lead to transcripts that carry premature termination codons and thus destabilized by the NMD pathway (Lewis, Green and Brenner, 2003). Alternative splicing coupled to NMD (AS-NMD) is an important component of the molecular circuitry essential for the generation of neuron-specific alternative splicing programs and neuronal differentiation (Makeyev *et al*., 2007). Colak and colleagues (2013) showed that NMD regulates Robo3.2 protein levels by inducing the degradation of *Robo3*.*2* transcripts in axons that encounter the floor plate (Colak *et al*., 2013). NMD pathway was shown to control GLUR1 surface levels through degradation of *Arc* and *Prkag3*, thus regulating synaptic plasticity. The authors also show that core components of the NMD machinery are localized to neurites (Notaras *et al*., 2019). In both studies alternative splicing led to intron retention in the 3’ UTR resulting in premature termination codons within their reading frame. Our findings suggest that SNRNP70 could be a critical player in AS-NMD regulation of gene expression outputs in developing neural circuits. This mechanism would be particularly useful during axon guidance and synaptogenesis where rapid changes in protein expression are required for the axonal growth cone and focal axonal areas to rapidly responds to signals from the environment.

In conclusion, our studies reveal unexpected cytoplasmic functions of a key spliceosomal factor, SNRNP70, during the development of neuronal connections. Further work will be needed to examine the full extent of the biological functions of SNRNP70 in neurites and the mechanisms by which it controls the local transcriptome composition and influences the establishment and maintenance of neuronal connectivity.

## Acknowledgements

We thank Prof. David Stanek for kindly giving us the human SNRNP70 construct, Prof. Martin Meyer for the HuC:Gal4 construct, Triona Fielding for technical support throughout the work, and Dr Rachel Moore and Prof. Jon Clarke for assistance in using the Zeiss Airyscan confocal imaging system. We also thank the King’s College London Guy’s campus fish facility staff for their fish husbandry and care. This study was supported by the Biotechnology and Biological Sciences Research Council (BB/P001599/1 to C.H. & BB/M007103/1 and BB/R001049/1 to E.V.M.) and the Wellcome Trust (Tech Dev. WT093389 to C.H.).

## Author contributions

C.H. conceived the research project and secured funding. N.N., P.M.G., F.H., R.T., E.V.M., and C.H. designed the experiments. N.N., P.M.G. and R.T. conducted the experiments. N.N. analyzed data. F.H. performed the bioinformatic analysis of RNA sequencing data. N.N. wrote the manuscript with input from all authors.

## METHODS

### Zebrafish lines and husbandry

Zebrafish were reared at 28.5°C on a 14 hr light/10 hr dark cycle. Embryos produced by natural crosses were raised in Danieau’s solution (58 mM NaCl, 0.7 mM KCl, 0.4 mM MgSO4, 0.6 mM Ca(NO3)2, 5.0 mM HEPES, pH 7.2). We used the following existing transgenic lines and strains: *Tg(hb9:GFP)*^*ml2Tg*^ (Flanagan-Steet *et al*., 2005), *Tg(ubi:ERT2-Gal4)*^*nim10Tg*^ (Gerety *et al*., 2013) and AB. The following lines were newly generated for the purpose of this study: *snrnp70*^*kg163*^, *Tg(UAS:hSNRNP70-eGFP)*^*kg323Tg*^ and *Tg(UAS:hSNRNP70ΔNLS3xNES-eGFP)*^*kg325Tg*^. This work was approved by the local Animal Care and Use Committee (King’s College London), and was carried out in accordance with the Animals (Experimental Procedures) Act, 1986, under license from the United Kingdom Home Office.

### Generation of *snrnp70* knock-out allele

The *kg163* null allele of *snrnp70* was generated by injecting the one-cell stage AB embryos with 250 pg Cas9 mRNA and 40 pg sgRNA. The sgRNA was generated by inserting the guide sequence GAACGGTATGGGGTCCCGC into vector pDR274 (Addgene) and transcribing using the T7 RiboMAX RNA Production kit (Promega-P1320). Cas9 was transcribed from the plasmid pCS2-Cas9 (Addgene) using the SP6 mMessage mMachine kit (Ambion-AM1340). Adult F0 animals were outcrossed with AB zebrafish and F1 offspring embryos were screened for CRISPR-induced ‘indel’ mutations using high melt resolution (HRM) analysis (HRM primers in Table S6). Embryos with shifted HRM curves compared to un-injected AB embryos were sequenced (genotyping primers in Table S6) to reveal the type of mutation. This led to the identification of an F0 founder that produces embryos with a 13bp deletion and was used to generate the stable lines used in this study.

### Generation of transgenic constructs

Sequences for all primers used are listed in Table S6. The expression construct *pT2 5UAS MCS eGFP* construct was generated by digesting *pN2 5UAS eGFP* (Fredj *et al*., 2010) with AseI and AflII restriction endonucleases (NEB). The 1.4 kb product (containing a 5xUAS and a hsp70 minimal promoter, a multiple cloning site, the enhanced green fluorescent protein sequence and an SV40 polyA) was blunted at both ends using a Klenow fragment of DNA polymerase I (NEB) and cloned into the *EcoRV* site of the *pminiTol2* plasmid (ref). To generate the *pT2 5UAS hSNRNP70-eGFP* expression construct, a 1.4 kb fragment containing the *hSNRNP70* sequence was isolated from the *pN1 CMV hSNRNP70-eGFP* plasmid (Huranová *et al*., 2010) (gift from David Stanek, Institute of Molecular Genetics, Czech Republic) by digestion with NheI and BamHI restriction endonucleases (NEB). The fragment was then directionally cloned into the *NheI/BamHI* sites downstream of the *UAS* motifs of the *pT2 5UAS MCS eGFP* construct generating C-terminally fused enhanced green fluorescent protein (*hSNRNP70-eGFP*). To generate a cytosolic hSNRNP70 expression construct, the *pT2 5UAS hSNRNP70-eGFP* plasmid was used as a template for sequential site directed mutagenesis using QuickChange II (Agilent-200523) to mutate the two nuclear localization signals (NLS) found on the gene as previously described (Romac, Graff and Keene, 1994). Briefly, for NSL1 the hSNRNP70 amino-acid sequence 166-GKK-168 was changed to 166-DFP-168 and for NLS2 the amino-acid sequence 186-GWRPRR-191 was altered to 186-ANLVDL-191. To introduce three copies of MEKK nuclear export signals (NES; amino acid sequence: ALQKKLEELELDE) at the C-terminal end of hSNRNP70 protein, we performed two rounds of restriction free cloning using primers encoding the NES sequence. Following the second round of restriction free cloning, we identified clones that contained 2x as well as 3x copies of NES. We thus decided to use the latter one for all of our cytoplasmic SNRNP70 rescue experiments. All constructs generated were confirmed by nucleotide sequencing.

### DNA microinjections for sparse labelling and generation of transgenic lines

Fertilized zebrafish AB embryos were microinjected with 1.8 nl of an injection solution containing 15-25 ng/μl DNA and 17 ng/μl Tol2 transposase mRNA. For single-cell analysis of hSNRNP70 localization, the *pT2 5UAS hSNRNP70-eGFP* expression construct was co-injected with a *HuC:Gal4* plasmid (gift from Martin Meyer, KCL, UK) together with 100 μM UTP-Cy5 (Perkin Elmer-NEL583001EA) inside a single blastomere at 4-8 cell stage embryos. For establishing stable transgenic lines, animals injected with either *pT2 5UAS hSNRNP70-eGFP* or *pT2 5UAS hSNRNP70ΔNLS3xNES-eGFP* DNA constructs together with transposase mRNA were raised to adulthood. Adult F0 animals were outcrossed with *Tg(ubi:ERT2-Gal4)* zebrafish and F1 offspring were screened for germline transmission of the fluorescent transgene.

### Time-lapse imaging of cytoplasmic SNRNP70

Embryos injected with *pT2 5UAS hSNRNP70-eGFP* and *HuC:Gal4* DNA plasmids together with UTP-Cy5 were mounted in 1% low melting point agarose (Sigma) in Danieau’s solution. Imaging was performed using a Zeiss LSM 880 Fast Airyscan confocal microscope equipped with 2x GaAsP spectral detectors and a 20x/1.0 NA water-immersion objective (Carl Zeiss). Excitation was provided using 488 nm (for GFP) and 633 nm (for Cy5) solid state lasers. High resolution images were captured at 0.2-0.33 Hz at a 0.06 × 0.06 μm resolution (1172 × 1172 pixel) and 1 A.U. pinhole aperture and then processed for Airyscan imaging.

### 4-OHT treatment for GAL4 induction

4-Hydroxytamoxifen (4-OHT; Abcam-ab141943) was dissolved in ethanol at a final stock concentration of 50 mM and stored in the dark at –20°C. To induce Gal4 activity in Gal4ERT2-expressing offspring, embryos at mid-blastula stage (∼ 4 hpf) were washed in a petri dish containing Danieau’s solution, all medium was removed and replaced with fresh Danieau’s solution containing 2 μM of 4-OHT. The treated embryos were immediately put into a dark 28.5°C incubator to allow for efficient induction and remained in 4-OHT-containing Danieau’s solution for the rest of the experiment.

### Spinal motor axon mosaic experiment

Cell transplantations were done as previously described(Thomas-Jinu *et al*., 2017). Briefly, progenies from incrosses of *snrnp70*^*+/−*^;*Tg(hb9:GFP)* were used in this study. Around 20 dorso-posterior epiblast cells collected from late gastrula stage donor embryos (80%-100% epiboly) were placed at the equivalent location in the same stage wild-type host embryo. The donor embryos were identified as siblings or nulls by PCR at 28 hpf. Embryos that led to the incorporation of the donor cells in the ventral spinal cord and GFP-positive motor axons were selected for further analysis.

### Tissue preparation and cryosectioning for histology

Zebrafish embryos and larvae (48 – 120 hpf) were euthanized with 4 mg/mL Tricaine (MS-222; Sigma-A5040) and immersion fixed overnight at 4°C in 4% paraformaldehyde. Fixed animals were embedded in an agar-sucrose solution (1.5% agar, 5% sucrose) in PBS and let to set. The agar blocks were taken out of their moulds, placed into 30% sucrose solution. Following an overnight incubation at 4°C, blocks are frozen by gradual immersion into liquid nitrogen and stored at −80°C until sectioning. Transverse sections of 20 μm thickness were cut using a Bright Instruments-OTF5000 cryostat and subsequently stored at −80°C until further use.

### Immunohistochemistry

For whole mount immunohistochemistry, 4% paraformaldehyde-fixed embryos were washed with PBS, permeabilized with 0.25% Trypsin in PBS for 20 minutes and blocked with 10% goat serum/PBS at room temperature (RT) for 1 hour. Embryos were incubated with anti-GFP 1:500 (Torrey Pines Biolabs-TP401), anti-SNRNP70 1:100 (Sigma-AV40276; Lot: QC9623), anti-SV2 1:200 (DSHB-AB2315387), anti-F59 1:10 (DSHB-AB528373), anti-F310 1:10 (DSHB-AB531863) diluted in 10% goat serum/PBST (1% Triton X-100) at 4°C overnight, washed with PBST (1% Triton X-100) and incubated with Alexa Fluor 488/568/633 secondary antibodies (1:500; Thermo Fisher Scientific) diluted in 10% goat serum/PBST (1% Triton X-100) at 4°C overnight. For staining Acetyl Choline receptors (AChRs) we incubated embryos with 1:100 α-Bungarotoxin (α-BTX)-555 conjugate (Thermo Fisher Scientific-B35451) at 4°C overnight. In some cases, HOECHT (Thermo Fisher Scientific-H3569) or DAPI (Sigma-D8417) were used at 1:1000 to stain cell nuclei. Cryosections were rehydrated in PBS for 10 minutes and then blocked in for 1 hour before incubating primary antibodies at 4°C overnight. Sections were then washed and incubated with secondary antibodies for 2 hours at room temperature.

### Imaging of immunofluorescent samples

Following immunohistochemistry, samples were imaged using a Zeiss Axio Imager.Z2 LSM 800 confocal microscope equipped with 2x GaAsP spectral detectors. Excitation was provided using 405, 488, 561 and 633 nm solid state lasers. Whole mount immunostainings of slow and fast muscle fibers using F59 and F310 antibodies, respectively, were imaged using a 20x/0.8 NA air objective (Carl Zeiss) at a 0.62 × 0.62 μm resolution (512 × 512 pixel) and 1 A.U with 1 μm sectioning z-interval. Whole mount labeling of pre- and posy-synaptic NMJ markers using anti-SV2 and α-BTX was imaged using a 40x/1.0 NA oil objective (Carl Zeiss) at a 0.31 × 0.31 μm resolution (512 × 512 pixel) and 1 A.U with 0.75 μm sectioning z-interval. Immunostainings of all other whole mount samples as well as cryosections were imaged using a 40x/1.0 NA oil objective (Carl Zeiss) at a 0.31 × 0.31 μm resolution (512 × 512 pixel) and 1 A.U. High magnification images of cryosections showing spinal cord and peripheral nerves were captured at a 0.08 × 0.08 μm resolution (512 × 512 pixels) and 1 A.U. pinhole aperture.

### Co-localization analysis of pre- and post-synaptic NMJ markers

NMJ analysis was done as previously described(Boon *et al*., 2009). Briefly, following confocal imaging of anti-SV2 and α-BTX within somites 10 and 11, datasets were opened in ImageJ. Anti-SV2 and α-BTX channels were merged and a maximum projection of a z-stack covering a depth of 15 μm was made. Only the ventral part of the myotomes was used for this analysis. Channels were split and background was subtracted using a rolling ball radius of 10 pixels. Finally, the overlap coefficient was calculated using the JACoP plugin.

### Zebrafish primary cell culture

Embryos of the *Tg(hb9:GFP)* line were collected, incubated overnight in Danieau’s solution containing 0.01% methylene blue, Penicillin-Streptomycin (Sigma-P4333) and 50 μg/mL Gentamicin (Gibco-15750060) and bleached at 28 hpf using 0.000017% sodium hypochlorite. Bleached embryos were then dechorionated using pronase (Sigma-10165921001), transferred into a tube containing 1 mL of cell dissociation solution (25% Cell Dissociation Buffer (enzyme free), 3.25×10^−3^ M EDTA, PBS, pH 8.0) and stored on ice. Embryos were then dissociated using a series of autoclaved Fisherbrand Pasteur pipets with cotton plugs with successively smaller bore diameters. After approximately 15 minutes, when chunks of embryos were no longer visible, the dissociated cells were passed through a Corning 40 μm cell strainer into a 50 mL centrifuge tube. Extra cell dissociation solution was used to rinse the cell strainer membrane, pushing through any material stuck on top. The tube was centrifuged at 300g for 7 minutes at 4°C. Cell dissociation solution was removed from above the cellular pellet, before resuspension in Zebrafish Neural Medium (ZNM) consisting of Leibovitz L-15 medium (Gibco-11415049), 1X N-2 Supplement (Gibco-17502048), 1X MACS NeuroBrew-21 Supplement, 10ng/mL BDNF (Sigma-B3795), 2% FBS, 1X Penicillin-Streptomycin (Sigma-P4333) & 50 μg/mL Gentamicin (Gibco-15750060). Cell solution was subsequently divided, plating onto Corning BioCoat Poly-D-Lysine/Laminin-coated coverslips in a 24-well-plate. The plate was sealed by wrapping with parafilm and left overnight at RT. Cells were fixed at DIV1. ZNM was removed from each well before rinsing once with PBS. 1 mL of 4% PFA was added to each well for 30 minutes at RT. Following removal of PFA, coverslips were washed 2 × 5-minutes with PBST (1%Triton X-100). Coverslips were then incubated with blocking solution (PBST and 5% Goat Serum) for 1 hour at RT. This was removed and coverslips were incubated with fresh blocking solution containing anti-GFP 1:1000 (Torrey Pines Biolabs-TP401) and anti-SNRNP70 1:200 (Sigma-AV40276) overnight at 4°C. This solution was removed, and coverslips subsequently underwent 3X PBST washes at RT whilst on a shaker. They then had further blocking solution added to them containing Alexa Fluor 488/568 secondary antibodies (1:1000; Thermo Fisher Scientific) in addition to DAPI (1:2000; Sigma-D8417). Coverslips were incubated at RT for 2 hours. After removing the solution from coverslips, they underwent 3 X PBST washes over the course of a few hours. Coverslips were mounted onto glass slides with 10μl Calbiochem FluorSave Reagent. Samples were imaged using a Zeiss Axio Imager.Z2 LSM 800 confocal microscope equipped with 2x GaAsP spectral detectors and a 40x/1.3 NA oil objective (Carl Zeiss). Excitation was 405 nm (for DAPI), 488 nm (for GFP) and 561 nm (for SNRNP70). Images were captured at a 0.2 × 0.2 μm resolution (512 × 512 pixel) and 1 A.U. pinhole aperture.

### Morphological analysis of motor nerves

Live *Tg(hb9:GFP)* embryos were mounted in 1% low melting point agarose (Sigma) in Danieau’s solution. Imaging was performed using a Zeiss Axio Examiner.Z1 LSM 800 confocal microscope equipped with 2x GaAsP spectral detectors and a 20x/1.0 NA water-immersion objective (Carl Zeiss). Excitation was provided using a 488 nm solid state laser. Image stacks of motor nerves were captured at a 0.62 × 0.62 μm resolution (512 × 512 pixel) and 1 A.U. pinhole aperture with 0.5 μm sectioning z-interval. Datasets were opened in ImageJ and motor nerves in somites 9-12 were traced using the Simple_Neurite_Tracer plugin. Motor nerve length was measured from the spinal cord exit point to the most distal end of the nerve. The length of myosepta innervation was measured from the most ventral part of the motor nerve until the distal end of the nerve near the horizontal myoseptum and normalised to the total distance to the horizontal myoseptum to get the percentage of myosepta length innervated. Motor nerve thickness was quantified by making maximum intensity projections and using a straight line to measure the thickness of the nerve at the level of the choice point at the horizontal myoseptum.

### Assessment of startle response

Touch-evoked startle response was measured as previously described (Telfer *et al*., 2010). Briefly, startle responses were elicited by gently touching 48 hpf animals using a blunt tool and measured using a scale from 0 to 3 as follow: 0, no movement; 1, flicker of movement but no swimming; 2, movement away from probe but with impaired swimming; and 3, normal swimming.

### RNA sequencing

Embryos derived from an outcross of *snrnp70*^*+/−*^;*Tg(ubi:ERT2-Gal4);Tg(UAS:hSNRNP70ΔNLS3xNES-eGFP)* with *snrnp70*^*+/−*^;*Tg(ubi:ERT2-Gal4)* animals were treated with 2 μM 4-OHT (as described above). Treated embryos were screened for GFP fluorescence and also sorted into siblings and nulls (based on the brain morphogenesis phenotype) at 24 hpf to produce the following four experimental groups: sib/GFP^−^, sib/GFP^+^, null/GFP^−^ and null/GFP^+^. Total RNA was isolated from these four pools of embryos at 28 hpf using a RNeasy Mini kit (QIAGEN-74104) and eluted in nuclease-free water. Total RNA samples (100-500 ng) were hybridized with Ribo-Zero Gold to substantially deplete cytoplasmic and mitochondrial rRNA from the samples. Stranded RNA sequencing libraries were prepared as described using the TruSeq Stranded Total RNA Library Prep Gold kit (Illumina-20020598) with TruSeq RNA UD Indexes (Illumina-20022371). Purified libraries were qualified on an Agilent Technologies 2200 TapeStation using a D1000 ScreenTape assay (Agilent-50675582 and 50675583). The molarity of adapter-modified molecules was defined by quantitative PCR using the Kapa Biosystems Kapa Library Quant Kit (Roche-KK4824). Individual libraries were normalized to 1.30 nM in preparation for Illumina sequence analysis. Sequencing libraries (1.3 nM) were chemically denatured and applied to an Illumina NovaSeq flow cell using the NovaSeq XP chemistry workflow (Illumina-20021664). Following transfer of the flowcell to an Illumina NovaSeq instrument, sequencing was performed using a NovaSeq S1 reagent Kit (Illunima-20027465) to obtain 50 nucleotide paired-end reads. All library preparation and sequencing steps were carried out by the Huntsman Cancer Institute High-Throughput Genomics facility, University of Utah, USA.

### Bioinformatics

For gene expression analyses and splicing analyses of 28 hpf RNA sequencing data, reads were aligned to zebrafish VASTDB library using VAST-TOOLS alignment tool(Tapial *et al*., 2017) and gene read counts output was enabled. Alignment was done as follows:

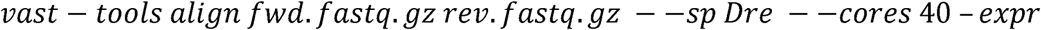

Splicing data from each sample were combined using VAST-TOOLS’ combine tool to obtain an INCLUSION table. Differential splicing analysis between samples was then determined using VAST-TOOLS’ “diff” command and a minimum change in Percent Spliced In (dPSI) value of 0.05 was requested. This was done as follows:

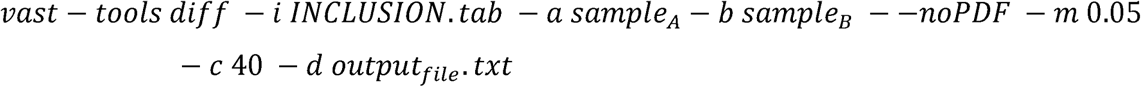

Differential gene expression analysis was carried out using edgeR package with the estimateGLMRobustDisp function (Robinson et al, 2010). Downstream processing of splicing and gene expression data were done in R using custom scripts.

### NMD prediction

For the prediction of NMD sensitivities upon splicing change, custom transcript annotations with alternatively spliced segments were generated using BioConductor packages. Coding sequences for each transcript were built and subsequently predicted for NMD-inducing features using in-house R scripts.

### GO enrichment analysis

GO term analysis of the affected groups of genes was performed using the Gene Ontology Enrichment resource (Ashburner *et al*., 2000; Carbon *et al*., 2019). We identified significantly enriched terms (*P* < 0.05) using the Panther overrepresentation test for enrichment(Mi *et al*., 2019) with a Fisher’s exact test and used the following databases: GO/Biological processes, GO/Cellular component and GO/Molecular function.

### Quantitative and semi-quantitative RT-PCR

Total RNA was extracted as described above for RNA sequencing. cDNA was synthesized using a First strand cDNA synthesis kit (Thermo Fisher Scientific-K1612) after treating the RNA isolated with DNaseI (QIAGEN-79254). Semi-quantitative RT-PCR reactions were performed using DreamTaq Green PCR Master Mix (Thermo Fisher Scientific-K1081) on an Eppendorf thermocycler. RT-qPCR reactions were performed using LightCycler 480 SYBR Green I Master kit (Roche-04707516001) on a LightCycler 96 system (Roche) and data analyzed using the LightCycler96 software. Expression levels of transcripts were first normalized to an endogenous control gene *actin-b1*, to obtain ΔCt values and then normalised again to the total expression of the endogenous gene to obtain ΔΔCt values. All primer sequences used are listed in Table S6.

### Identification of *z+agrn* micro-exons

Through sequence alignment of zebrafish *agrn* gene with mouse and human *z+agrn* isoforms we mapped the two putative micro-exons between exons 33 and 34 of the zebrafish AGRN locus. We performed RT-PCR using a forward primer in exon 33 and a reverse primer in exon 34 followed by DNA sequencing of bands obtained.

### Statistical analyses

The number of embryos and samples (n) and definition of statistical significance are indicated in the figure legends. Unpaired t-tests were used to compare means. For multiple comparisons, a One-way ANOVA test was used. For association analyses in gene expression and alternative splicing groups, a Fisher’s exact test was used. The criterion for statistical significance was set at *P* < 0.05 and results are represented as mean ± SEM. Mean values were calculated and plotted using GraphPad Prism (GraphPad Software).

## SUPPLEMENTARY DATA LEGENDS

**Figure S1: Characterization of the *in vivo* localization of SNRNP70**

**(A)** Single plane confocal images of cryosections derived from *Tg(hb9:GFP)* animals at 48 hpf and 120 hpf stained with anti-SNRNP70, anti-GFP and HOECHST to mark nuclei. Right: insets from the regions indicated on the left depicting nuclear (arrowheads) and axonal (arrows) localization of SNRNP70 in motor neurons. Representative images from two independent experiments. Scale bar, 25 μm.

**(B)** Single plane confocal image of cryosection derived from a wild-type animal at 120 hpf stained with anti-SNRNP70, anti-SV2 and α-BTX. Right: insets from the regions indicated on the left showing SNRNP70 localization at neuromuscular junctions (NMJs), whose identification was based on the apposition of SV2-labelled motor neurons and α-BTX-positive AChRs on muscle fibers (arrows). Representative image from two independent experiments. Scale bar, 25 μm.

**Figure S2: Generation of a *snrnp70* knock-out allele**

**(A)** Genomic architecture of *snrnp70* locus. The *cas9* guide RNA/Cas9 nuclease target is shown within exon2 which resulted in a *snrnp70* mutant allele (*kg163*) that has a 13 base-pairs deletion.

**(B)** Schematic diagram indicating the protein domains of the WT SNRNP70 protein.

**(C)** Protein sequences of WT and *kg163* mutant allele. The sequences are color-coded according to the protein domains in **(B)** which was the result of alignment with mouse and human SNRNP70 sequences. In *kg163* allele, there is a frameshift after amino-acid 16 and a termination codon 66 amino-acids later resulting in a truncated protein.

**(D)** Single plane confocal images at the level of the spinal cord from sibling and null embryos at 24 hpf stained with anti-SNRNP70 and HOECHST to mark nuclei. Representative images from two independent experiments. Scale bar, 15 μm.

**(E)** Representative images of sibling and null embryos at 24 hpf showing the gross morphology of the animals at this stage. Arrows indicate brain morphogenesis defects such as smaller ventricles and neuronal cell death in the anterior CNS of null animals.

**Figure S3: Motor neuron growth and synaptogenesis following the loss of SNRNP70**

**(A)** Representative confocal images of *Tg(hb9:GFP)* sibling and null embryos at 48 hpf showing motor nerves innervating the myotome. Green arrows indicate abnormal branching within the myotome, blue arrows point towards aberrant branching and magenta arrows show reduced innervation of the lateral myosepta. Scale bar, 50 μm.

**(B-C)** Quantifications showing the thickness of motor nerves and the percentage of lateral myosepta innervated in the two groups. All graphs show mean values ± SEM. **** *P* < 0.0001, two-tailed unpaired t-test, *n* = 10 animals per group in two independent experiments.

**(D)** Confocal images of sibling and null embryos at 48 hpf stained with anti-SV2 antibody to mark the pre-synaptic locations and a fluorescently tagged α-BTX to label post-synaptic structures. Scale bar, 25 μm.

**(E)** Quantification of the degree of overlap between SV2 and α-BTX. The graph shows mean values ± SEM. **** *P* < 0.0001, two-tailed unpaired t-test, *n* = 18 animals per group in three experiments.

**(F-G)** Confocal images of *Tg(hb9:GFP)* sibling and null embryos at 28 hpf stained with anti-F59 and anti-F310 to mark slow and fast muscle fibers, respectively. Representative images from two independent experiments. Scale bar, 50 μm.

**Figure S4: Overexpression of full-length hSNRNP70-eGFP restores normal development**

**(A)** Schematic diagram depicting the tagged construct used to generate the hSNRNP70-eGFP transgenic line. The two endogenous nuclear localization signals (NLS) are also shown. Below: A single plane confocal image within the spinal cord of *Tg(ubi:ERT2-Gal4;UAS:hSNRNP70-eGFP)* embryos at 24 hpf showing eGFP fluorescence in all cells. Scale bar, 20 μm.

**(B)** Schematic diagram depicting the tagged construct used to generate the cytosolic hSNRNP70-eGFP transgenic line. The two endogenous nuclear localization signals (NLS) were mutated and three copies of nuclear export signals (NES) were also added. Below: A single plane confocal image within the spinal cord of *Tg(ubi:ERT2-Gal4;UAS:hSNRNP70ΔNLS3xNES-eGFP)* embryos at 24 hpf showing eGFP fluorescence that is restricted to the cytoplasm of all cells. Scale bar, 20 μm.

**(C)** Overexpression of hSNRNP70-eGFP in *Tg(ubi:ERT2-Gal4;UAS:hSNRNP70-eGFP)* embryos at 48 hpf. Embryos derived from the same clutch were sorted into four groups according to genotype: (1) sib/GFP^−^, (2) sib/GFP^+^, (3) null/GFP^−^ and (4) null/GFP^+^. Representative images from three independent experiments.

**(D)** Quantification of touch-evoked startle response at 48 hpf following overexpression of hSNRNP70-eGFP. The graph shows mean values ± SEM. **** *P* < 0.0001, One-way ANOVA, *n* = 12 animals per group in three independent experiments.

**(E)** Confocal images of embryos at 48 hpf following overexpression of hSNRNP70-eGFP. Embryos were stained with anti-SV2 antibody to mark the pre-synaptic locations and a fluorescently tagged α-BTX to label post-synaptic structures. Representative images from three independent experiments. Scale bar, 25 μm.

**(F)** Quantification of the degree of overlap between SV2 and α-BTX at 48 hpf following overexpression of hSNRNP70-eGFP. The graph shows mean values ± SEM. **** *P* < 0.0001; ** *P* < 0.01, One-way ANOVA, *n* = 12 animals per group in three independent experiments.

**Figure S5: RNA-seq analysis reveals changes in gene expression between sibling and null animals**

Data are based on RNA-seq in 28 hpf embryos derived from overexpression of cytosolic hSNRNP70-eGFP in sibling and null animals using the *Tg(ubi:ERT2-Gal4;UAS:hSNRNP70ΔNLS3xNES-eGFP)* line. Embryos were sorted into four groups according to genotype: (1) sib/GFP^−^, (2) sib/GFP^+^, (3) null/GFP^−^ and (4) null/GFP^+^.

**(A)** Plot showing a principal component analysis (PCA) of three biological samples per experimental group.

**(B)** MA plot illustrating gene expression changes between null/GFP^−^ vs sib/GFP^−^ based on a *P* value < 0.05. Red and blue dots represent upregulated and downregulated genes, respectively.

**(C)** MA plot illustrating gene expression changes between null/GFP^+^ vs null/GFP^−^ based on a *P* value < 0.05. Red and blue dots represent upregulated and downregulated genes, respectively.

**(D)** Four-way plot comparing gene expression changes between null/GFP^−^ vs sib/GFP^−^ and null/GFP^+^ vs null/GFP^−^. The 347 genes rescued by cytoplasmic SNRNP70 are shown in cyan while the 662 non-rescued genes are shown in magenta (threshold: *P* value < 0.05 for both comparisons). Within the non-rescued group there were cases where expression of the transgene aggravated the null effect as well as cases where the gene expression was unaffected in null/GFP^−^ but changed significantly in null/GFP^+^ animals.

**(E)** GO terms of the 347 genes rescued by cytoplasmic SNRNP70 from **(D)**, showing the fold enrichment in biological processes, cellular component and molecular function.

**Figure S6: GO term enrichment of pre-mRNA alternative splicing events rescued by cytoplasmic SNRMP70**

**(A)** Graphs of the GO terms of proteins whose pre-mRNA splicing was restored by cytoplasmic SNRNP70 showing selected fold enrichment in biological processes, cellular compartment and molecular function. Data are based on RNA-seq in 28 hpf embryos derived from overexpression of cytosolic hSNRNP70-eGFP in sibling and null animals using the *Tg(ubi:ERT2-Gal4;UAS:hSNRNP70ΔNLS3xNES-eGFP)* line. All 93 rescued alternative splicing events based on comparison between null/GFP^−^ vs sib/GFP^−^ and then between null/GFP^+^ vs null/GFP^−^ were used in this analysis.

**Figure S7: Alternative splicing events leading to NMD among the SNRNP70-regulated groups**

Data are based on RNA-seq in 28 hpf embryos derived from overexpression of cytosolic hSNRNP70-eGFP in sibling and null animals using the *Tg(ubi:ERT2-Gal4;UAS:hSNRNP70ΔNLS3xNES-eGFP)* line.

**(A)** Venn diagram showing the 2498 genes downregulated after loss of SNRNP70 (dark gray circle) and the 571 genes upregulated by cytoplasmic SNRNP70 (light gray circle). The intersection contains the 208 genes whose expression was significantly restored to control levels.

**(B)** Venn diagram showing the 2015 genes upregulated after loss of SNRNP70 (dark gray circle) and the 438 genes downregulated by cytoplasmic SNRNP70 (light gray circle). The intersection contains the 139 genes whose expression was significantly restored to control levels.

**(C)** Graph showing the percentage of genes undergoing significant alternative splicing within the gene expression groups shown. *** *P* < 0.001; ns, not significant, Fisher’s exact test.

**(D)** Graph showing the fraction of NMD-causing splicing events in genes from **(C)** that undergo alternative splicing. ** *P* < 0.01; ns, not significant, Fisher’s exact test.

**(E)** Graph showing the fraction of regulated splicing events leading to NMD among the splicing events either not rescue or significantly restored by cytoplasmic SNRNP70. * *P* < 0.05, Fisher’s exact test.

**Table S1: Gene expression changes following loss of SNRNP70**

List of genes whose expression was significantly increased or decreased following the loss of SNRNP70. The results are based on a comparison of cRPKM values between the null/GFP^−^ and sib/GFP^−^ groups. *P* < 0.05, build-in analysis from individual bioinformatics tool.

**Table S2: Rescued gene expression levels following cytoplasmic expression of SNRNP70**

List of genes whose expression levels were significantly restored following cytoplasmic expression of SNRNP70. The results are based on a comparison of cRPKM values between the null/GFP^−^ and sib/GFP^−^ groups and then between null/GFP^+^ and null/GFP^−^. *P* < 0.05, build-in analysis from individual bioinformatics tool.

**Table S3: Alternative splicing changes following loss of SNRNP70**

List of alternative splicing events significantly altered following the loss of SNRNP70. The results are based on a comparison of dPSI values between the null/GFP^−^ and sib/GFP^−^ groups. *MV*.*dPSI* > 0.1, build-in analysis from individual bioinformatics tool.

**Table S4: Rescued alternative splicing events following cytoplasmic expression of SNRNP70**

List of alternative splicing events significantly restored following cytoplasmic expression of SNRNP70. The results are based on a comparison of dPSI values between the null/GFP^−^ and sib/GFP^−^ groups and then between null/GFP^+^ and null/GFP^−^. *MV*.*dPSI* > 0.1, build-in analysis from individual bioinformatics tool.

**Table S5: Expression of spliceosomal components**

List of all genes encoding major and minor spliceosomal components and their gene expression levels in the form of cRPKM within each of the four experimental groups.

**Table S6: Primers used in this study**

List of all primers used for molecular biology, RT-PCR, RT-qPCR and genotyping *snrnp70*^*kg163*^ knock-out animals.

**Movie S1: SNRNP70 localizes in axonal RNP granules moving rapidly towards the neuronal cell body**

Time-lapse imaging of a zebrafish interneuron at 48 hpf expressing hSNRNP70-eGFP and UTP-Cy5 to mark RNA molecules inside RNPs. The SNRNP70^+^ panctum moves rapidly towards the neuronal cell body together with the RNP it is associated with.

**Movie S2: SNRNP70 localizes in cytoplasmic RNP granules slowly oscillating within the axonal compartment**

Time-lapse imaging of a zebrafish interneuron at 48 hpf expressing hSNRNP70-eGFP and UTP-Cy5 to mark RNA molecules inside RNPs. The SNRNP70^+^ panctum slowly oscillates within the axonal compartment together with the RNP it is associated with.

