## Supplementary figures and images for "Cytoplasmic pool of spliceosome protein SNRNP70 regulates the axonal transcriptome and development of motor connectivity"

### Figure S1

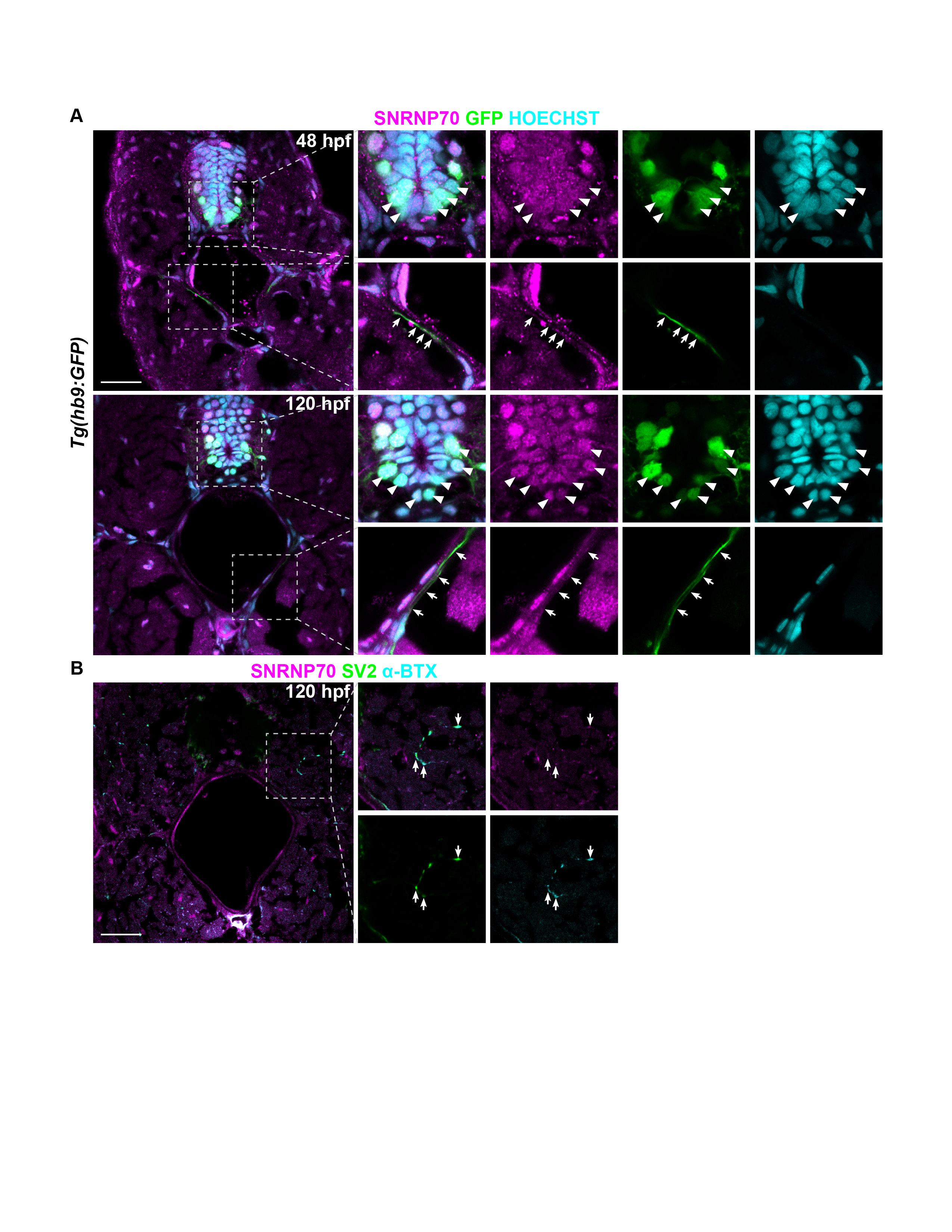

### Figure S2

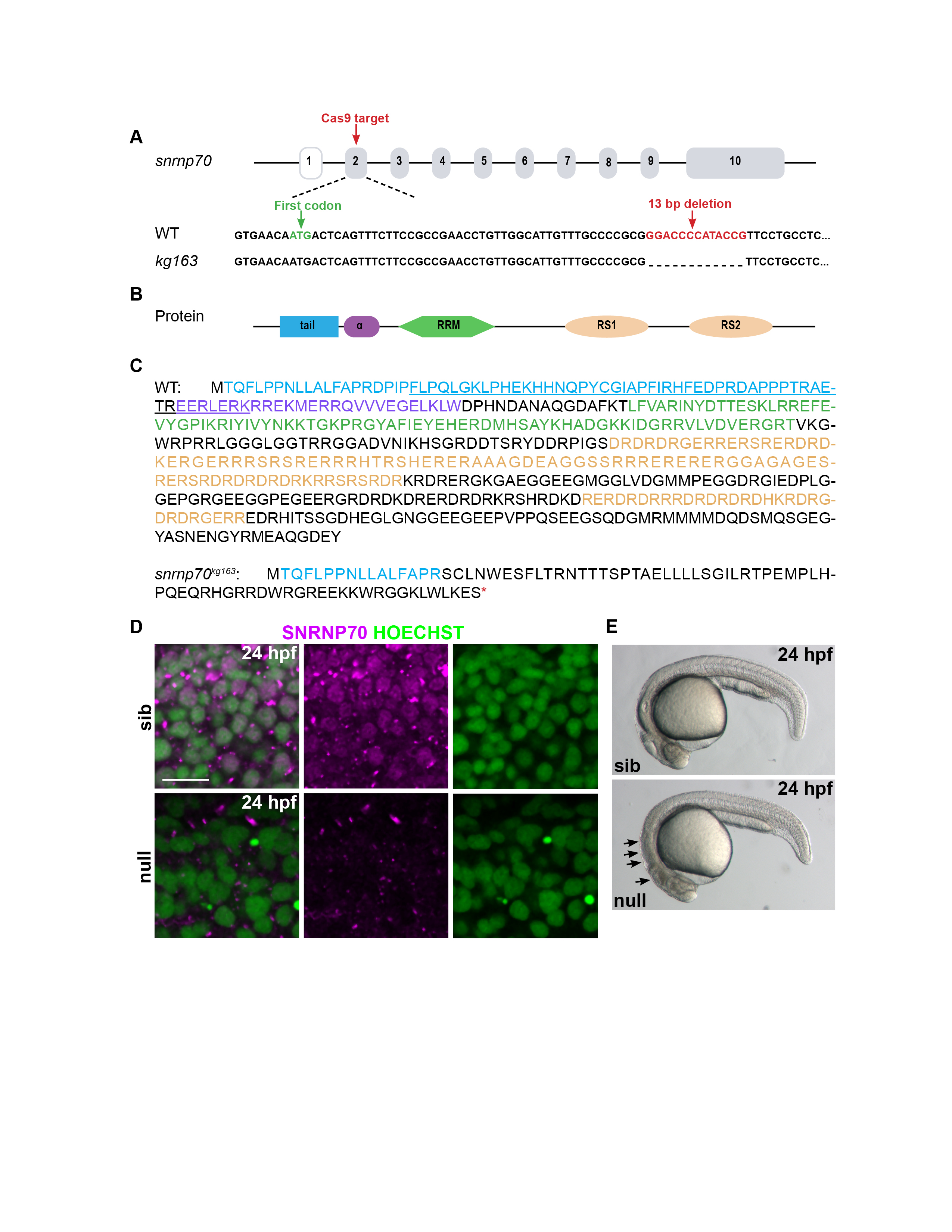

### Figure S3

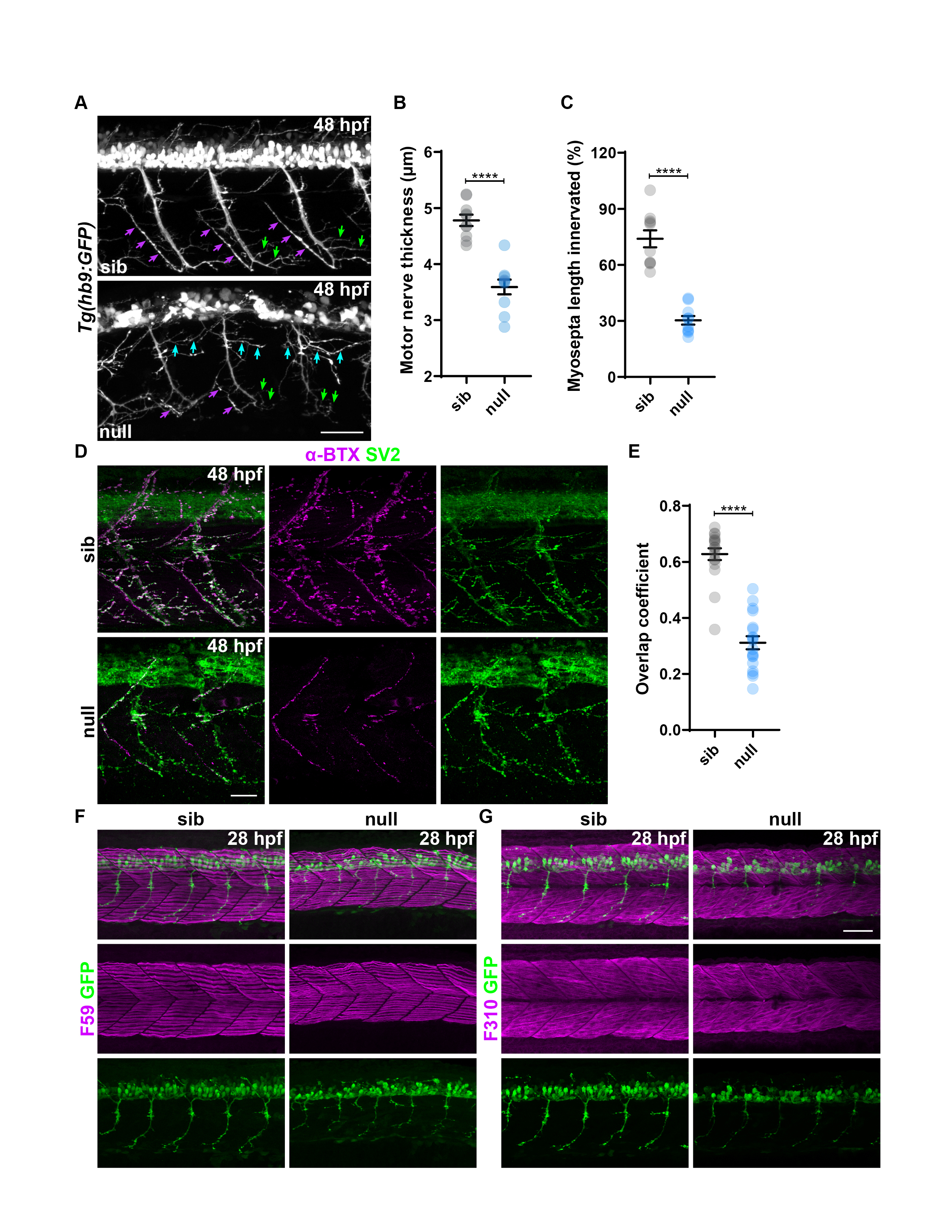

### Figure S4

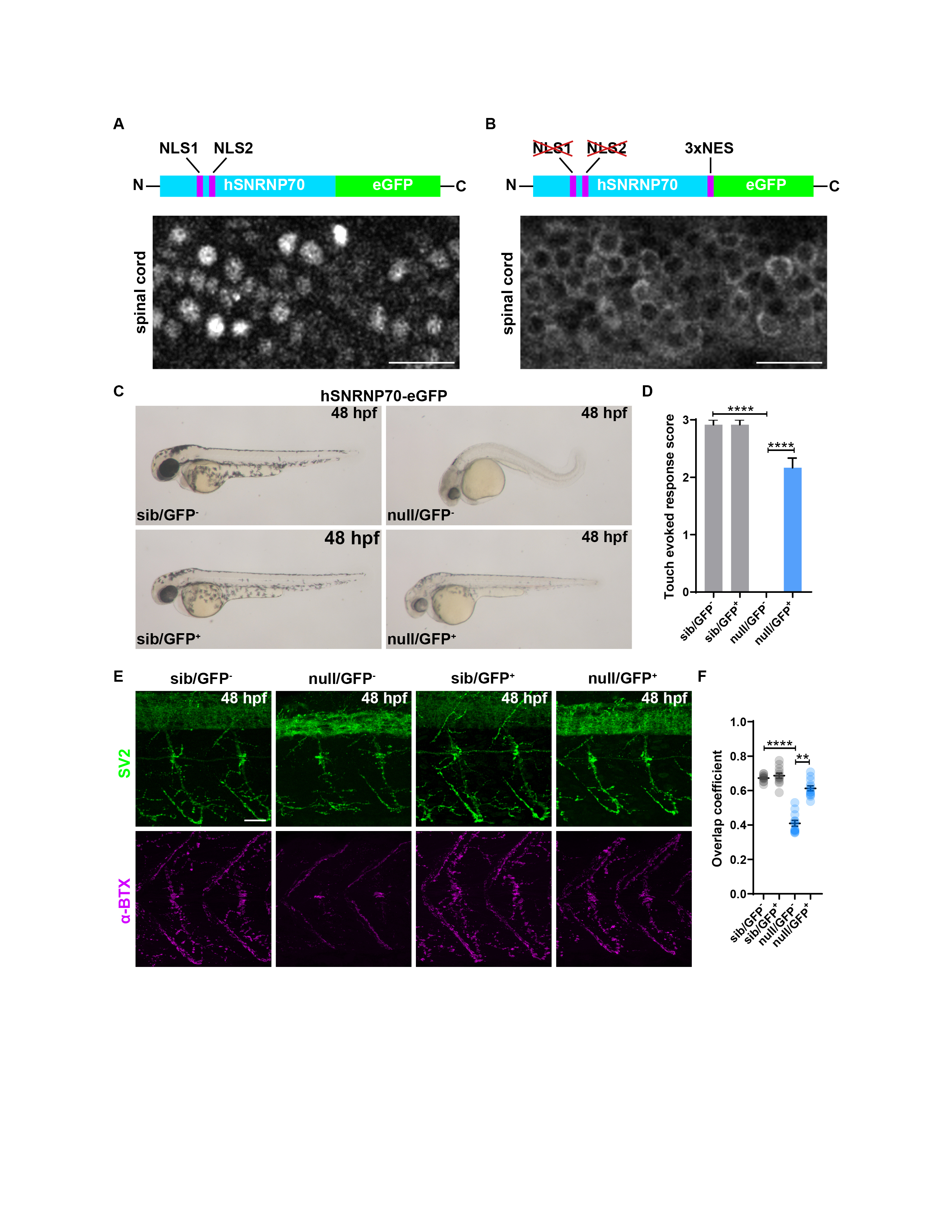

### Figure S5

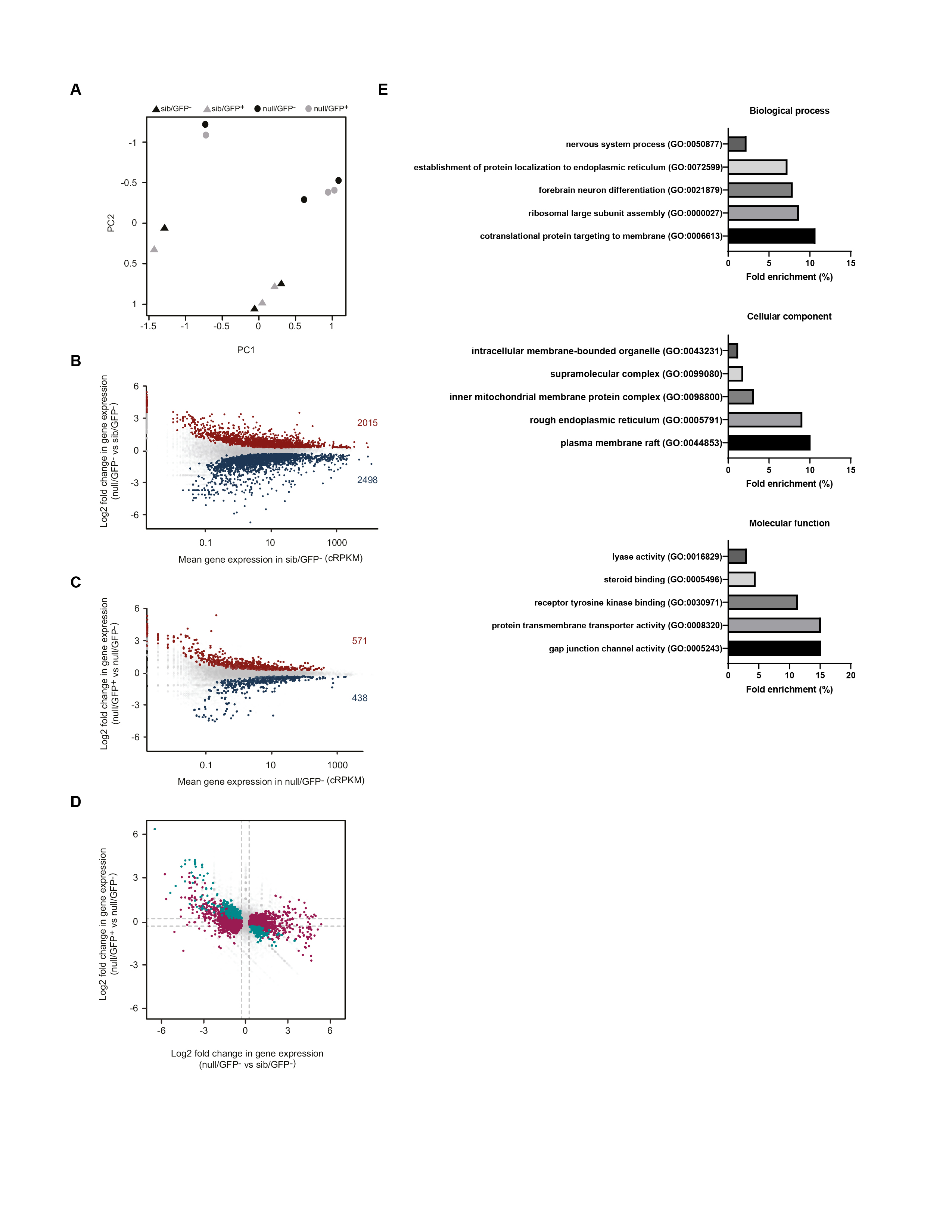

### Figure S6

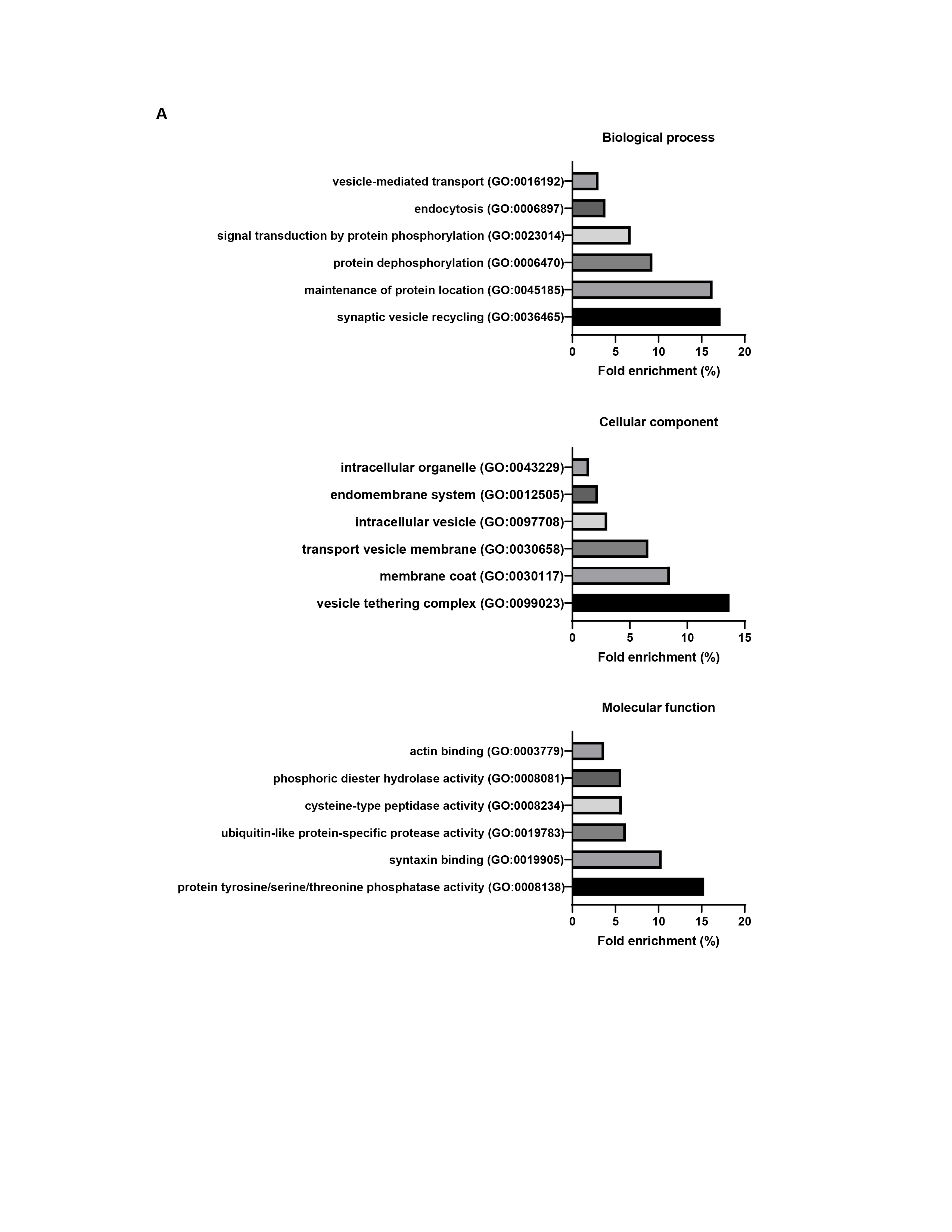

### Figure S7

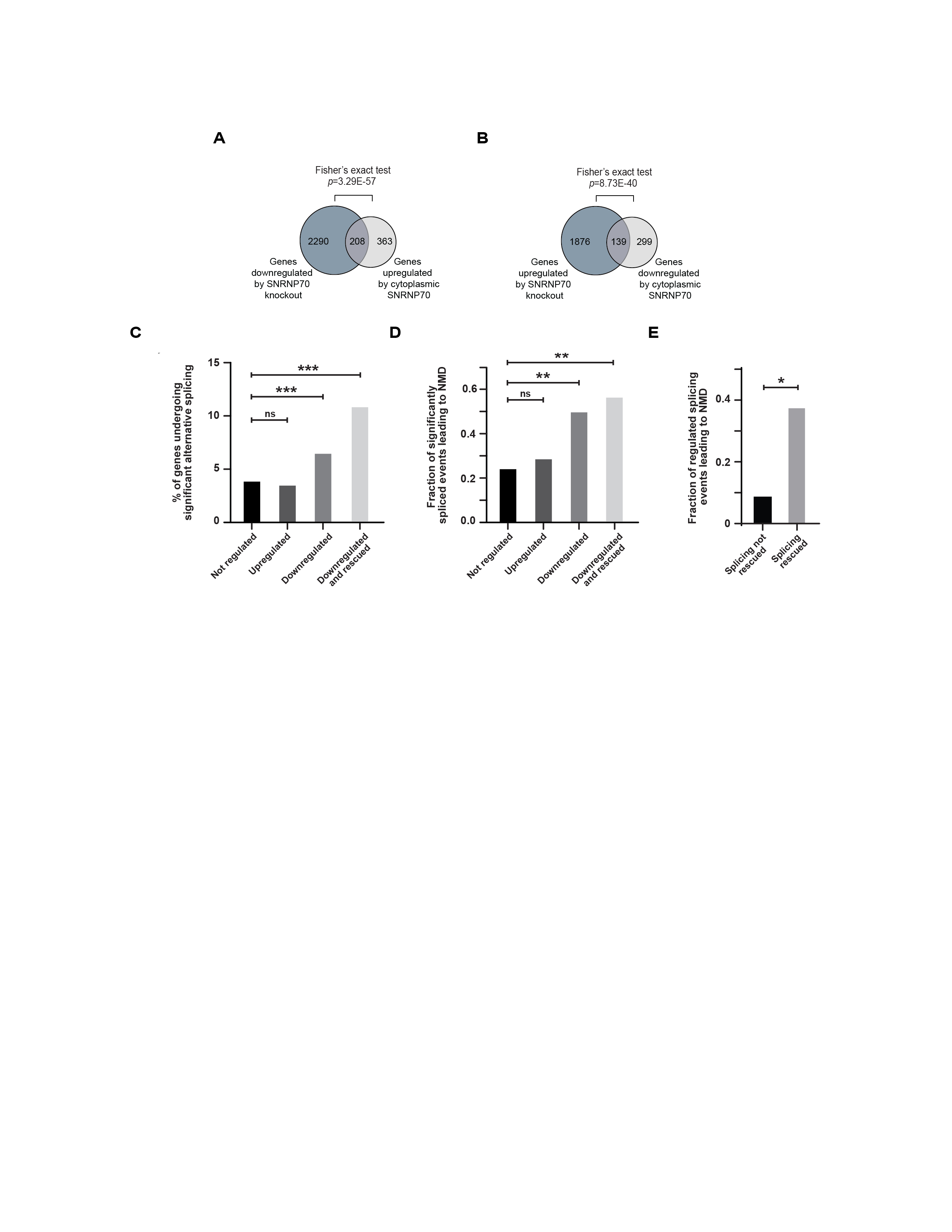
